# Avoidance engages dopaminergic punishment in *Drosophila*

**DOI:** 10.1101/2025.07.07.663268

**Authors:** Fatima Amin, Jasmine T. Stone, Christian König, Nino Mancini, Kazuma Murakami, Salil S. Bidaye, M.-Marcel Heim, David Owald, Utsab Majumder, Ilona C. Grunwald Kadow, Anna Pierzchlińska, Ashok Litwin-Kumar, Oliver Barnstedt, Bertram Gerber

## Abstract

It was classically suggested that behaviour can cause emotions (Darwin 1872). For example, smiling can make us feel happier, and in rodents the induced patterns of cardiac activity and breathing that are indicative of fear can in turn evoke it (Coles et al. 2022, Hsueh et al. 2023, Jhang et al. 2024). However, the adaptive significance of such feedback is unclear. We show that inducing backward movement, an element of avoidance behaviour in *Drosophila*, engages negative valence signals in these animals, and reveal the neuronal mechanisms and adaptive significance of this effect. We develop a paradigm with odours as conditioned stimuli paired with optogenetically induced backward movement instead of a punishing unconditioned stimulus, and combined these experiments with pharmacology, high-resolution video tracking, functional imaging, connectome analyses, and modelling. Our results show that backward movement engages dopaminergic punishment neurons and supports aversive memories. Such avoidance-to-punishment feedback counterbalances extinction learning and maintains learned avoidance, reducing the risk of further punishment. This can explain the long-standing “avoidance paradox”, the observation that avoidance adaptively persists even when it is successful and no punishment is received (Bolles 1972). Our results provide a neurobiologically grounded argument for an integrated view of behaviour organization and valence processing.

## Introduction

In his treatise on the relationship between emotions and behaviour, Charles Darwin suggested that not only can a particular emotional state engage a corresponding behaviour, but conversely we can adopt the emotional state corresponding to the behaviour we engage in (Darwin 1872; also see James 1884). This hypothesis sparked intense and ongoing debate, a prominent example of which is the question whether smiling can make us feel happier. After decades of controversy, a large-scale collaboration recently settled in favour of such facial feedback (Coles et al. 2022; also see Efthimiou et al. 2025). Likewise, in rodents induced increases in heart rate and changes in breathing pattern that are thought to be indicative of fear can in turn evoke it (Hsueh et al. 2023, Jhang et al. 2024). The function of such feedback to emotional states is unclear, however. We discovered that inducing backward movement engages negative valence signals in *Drosophila melanogaster*, and investigate the circuit mechanism and function of this effect.

*Drosophila* lends itself to such analyses as it permits the convenient expression of transgenes to induce, prevent, or measure neuronal activity during behaviour. This feature, combined with the numerical simplicity of the fly brain and the near-complete mapping of its connectome, has led to an understanding of the basic logic of how these animals learn to seek what is good and to avoid what is bad for them (Heisenberg 2003, Gerber & Aso 2017, Cognigni et al. 2018, Boto et al. 2020, Li et al. 2020, Modi et al. 2020, Davis 2023). Such learning takes place in the mushroom body, a higher brain centre that is evolutionarily conserved across insects. Its intrinsic neurons, called Kenyon cells (KCs), represent the sensory environment in a sparse and combinatorial manner, and are intersected by predominantly dopaminergic modulatory neurons. A subset of these mediate either negative or positive valence signals, evoked for example by electric shock punishment or sugar reward, which are conveyed to segregated compartments along the long, parallel axonal fibres of the KCs. Punishment and reward compartments feature output neurons that promote approach and avoidance, respectively. Upon coincidence of KC activity and a dopaminergic punishment signal, for example, the synapse from the KC onto the approach-promoting output neuron is depressed, shifting the balance across the output neurons as a whole to avoidance when the odour is encountered again. Despite the elegant simplicity of this logic, the connectome of the fly brain has revealed unexpected circuit complexity (Zheng et al. 2018, Li et al. 2020, Scheffer et al. 2020, Dorkenwald et al. 2024, Schlegel et al. 2024), suggesting corresponding behavioural and experiential complexity. We reasoned that this system should allow for a neurobiologically grounded understanding of the action-valence relationship, and asked whether avoidance can engage negative valence signals as reflected by the activity of dopaminergic punishment neurons.

The brain-descending “moonwalker” neurons elicit backward walking when experimentally activated (Fig. 1a; Extended Data Fig. 1) (Bidaye et al. 2014). We hypothesized that backward movement, an element of *Drosophilás* natural avoidance manoeuvres, might promote negative valence. We tested this idea in a modified Pavlovian conditioning paradigm using odours as conditioned stimuli and, unconventionally, the optogenetic induction of backward movement instead of a punishing unconditioned stimulus. These experiments are combined with behavioural pharmacology, high-resolution video tracking, functional imaging, connectomic analyses and a computational model. Our results suggest that avoidance engages dopaminergic punishment signals to maintain aversive memory, functionally counterbalancing extinction learning and reducing the likelihood of receiving further, potentially life-threatening, punishment. Such a mechanism can resolve the long-standing puzzle of why animals and humans alike often continue to avoid a cue predictive of punishment even when that avoidance is successful and no punishment is actually received (the “avoidance paradox”: Bolles 1972, LeDoux et al. 2017).

**Fig. 1.**
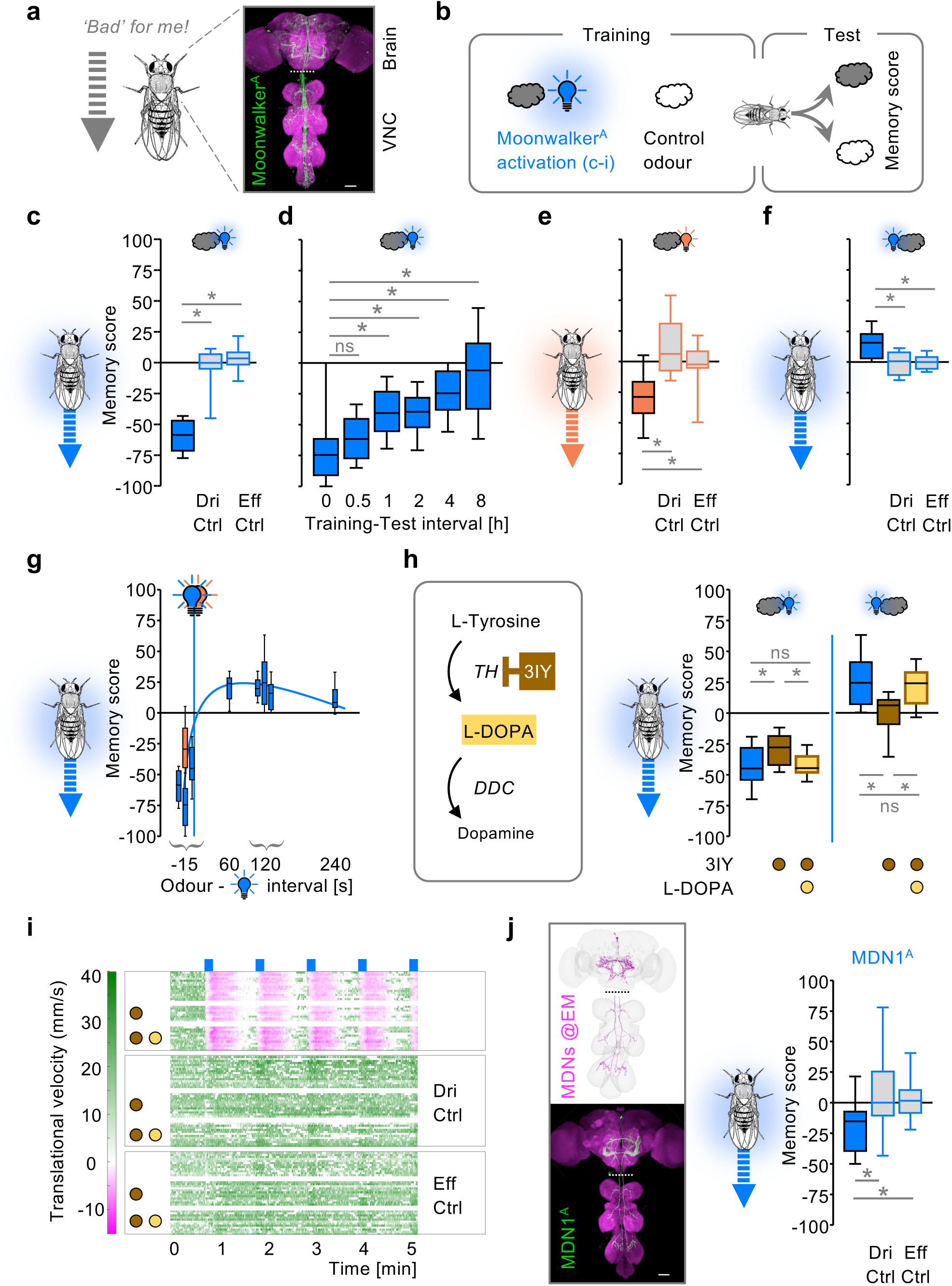
Moonwalker neuron activation engages a dopaminergic punishment signal. **a,** Proposed action-valence relationship and expression of the transgenic driver to induce backward movement. Moonwalker^A^>Chrimson^A^; magenta: neuropil labelling (anti-Bruchpilot), green: moonwalker neuron labelling (anti-GFP). VNC: ventral nerve cord. **b,** Overview of learning experiments. Clouds: odours. Light bulb: optogenetic activation of all moonwalker neurons (Moonwalker^A^) **(c-i)** or only the moonwalker descending neurons (MDN1^A^) **(j)**. **c,** Aversive memory by pairing odour with moonwalker neuron activation (blue: Moonwalker^A^>ChR2XXL^A^, grey: Driver control: Moonwalker^A^>+, Effector control: +>ChR2XXL^A^, +) (N= 20,18,18). **d,** As in **(c)**, for the indicated training-test intervals (N= 19,20,20,20,20,19). **e,** As in **(c)**, using Chrimson^A^ as effector (orange: Moonwalker^A^>Chrimson^A^, grey: Driver control: Moonwalker^A^>+, Effector control: +>Chrimson^A^) (N= 17,17,18). **f,** As in **(c)**, for a training procedure in which odour presentation followed moonwalker neuron activation (N= 18,15,15). **g,** Results for the experimental genotypes arranged according to the indicated intervals between odour and optogenetic activation (N= 23,23,23,18; 24; 24,24,24; 24) (includes data re-plotted from **(c-h)** for the-15 s and 120 s intervals). **h,** Inhibition of dopamine biosynthesis by 3-iodo-L-tyrosine (3IY), and its effects on learning from moonwalker activation (Moonwalker^A^>ChR2XXL^A^, intervals -15 s or 120 s; blue: control, brown: 3IY, light brown: additional supply of 3,4-dihydroxy-L-phenylalanine (L-DOPA)) (N= 23,24,23; 20,19,20). DDC: dopamine decarboxylase. TH: tyrosine hydroxylase. **i,** Analysis of movement upon the treatments in **(h)**. Shown is translational velocity (mm/s), colour-coded from magenta/backward to green/forward movement in relation to moonwalker activation (blue bars). Rows correspond to individual flies; the top three sets of rows show Moonwalker^A^>ChR2XXL^A^ flies; genetic controls, as in (c), are shown below (N= 12,8,12,16,12,12,12,12,12). **j,** EM reconstruction of the moonwalker descending neurons (MDNs, magenta) (grey mesh: brain and VNC) and expression of the transgenic driver covering them (MDN1^A^>Chrimson^A^, details as in **a**). Learning experiments as in **(c)**, showing aversive memory through MDN activation (blue: MDN1^A^>ChR2XXL^A^, grey: Driver control: MDN1^A^>+, Effector control: +>ChR2XXL^A^) (N= 21,17,17). Scale bars in **(a,j)**: 50 μm. Stippled lines in **(a,j)** indicate stitching of images of brain and VNC from the same animal, processed separately. Box-whisker plots show median, interquartile range (box) and 10th/90th percentiles (whiskers). Data were analysed across groups by Kruskal-Wallis tests (P< 0.05), followed by pairwise comparisons (Mann-Whitney U-tests, *P< 0.05 with Bonferroni-Holm correction) (ns: P> 0.05). Higher-resolution versions of **(a,j)** in Extended Data Fig. 1. NeuronIDs for **(j)** in Methods Supplemental Table 2. Additional information in Extended Data Fig. 2.

## Results

### Moonwalker neuron activation engages a dopaminergic punishment signal

To investigate action-valence relationships, we tested whether backward movement induced by moonwalker neuron activation can engage negative valence. Flies were trained in a differential conditioning task such that one odour was followed by optogenetic activation of the moonwalker neurons. A second, control odour was presented alone, followed by a choice test between the two odours (Fig. 1a,b). This revealed an aversive memory for the odour that had been associated with moonwalker neuron activation in the experimental genotype, but not in genetic controls treated the same (Fig. 1c).

Aversive moonwalker-memory subsided over 4-8 hours (Fig. 1d) and was observed also when using another optogenetic effector (Chrimson, rather than ChR2XXL: Fig. 1e). Interestingly, the punishing effect of moonwalker neuron activation has the same temporal ‘fingerprint’ as electric shock punishment. That is, whereas the presentation of an odour before an electric shock establishes strong aversive memory for the odour, presentation of an odour after the shock, at the moment of relief, leads to a characteristically weaker appetitive memory, a principle that applies across species (Gerber et al. 2019). We found that this general feature of punishment processing is also observed for moonwalker neuron activation (Fig. 1f,g).

As punishment is conveyed to the mushroom body KCs by dopaminergic neurons (Schwaerzel et al. 2003, Riemensperger et al. 2005, Claridge-Chang et al. 2009, Mao & Davis 2009, Aso et al. 2010, Aso et al. 2012), we hypothesized that learning from moonwalker activation is dopamine-dependent. Acutely supplementing the fly food with the drug 3IY, an inhibitor of the TH enzyme that is required for dopamine biosynthesis but is without effect on odour preference (Thoener et al. 2021), impaired punishment learning and abolished relief learning mediated by moonwalker neurons (Fig. 1h). Both these effects were rescued by additionally supplying L-DOPA (Fig. 1h). Critically, high-resolution video recordings showed that moonwalker-induced backward movement itself was unaffected by 3IY (Fig. 1i; Extended Data Fig. 2). These results show that moonwalker neuron activation both elicits backward movement and engages a dopaminergic punishment signal for associative learning.

The fly strain used for moonwalker neuron activation expresses the optogenetic effector relatively broadly (Bidaye et al. 2014) (Fig. 1a; Extended Data Fig. 1a). The covered neurons can be assigned to seven cell types, of which a single cell type comprising four neurons named moonwalker descending neurons (MDNs) is sufficient to induce backward movement (Bidaye et al. 2014). Using a fly strain to specifically activate these MDNs also produced a punishment memory when paired with odour (Fig. 1j; Extended Data Fig. 1b,c), allowing us to focus our subsequent analyses on these neurons. These analyses will reveal a reciprocal interaction between the MDNs and the flies’ olfactory memory centre, the mushroom body. To understand this reciprocal interaction, we first focus on how the mushroom body is connected to the MDNs (Fig. 2) and then investigate how the MDNs in turn affect mushroom body processing (Fig. 3-5).

**Fig. 2.**
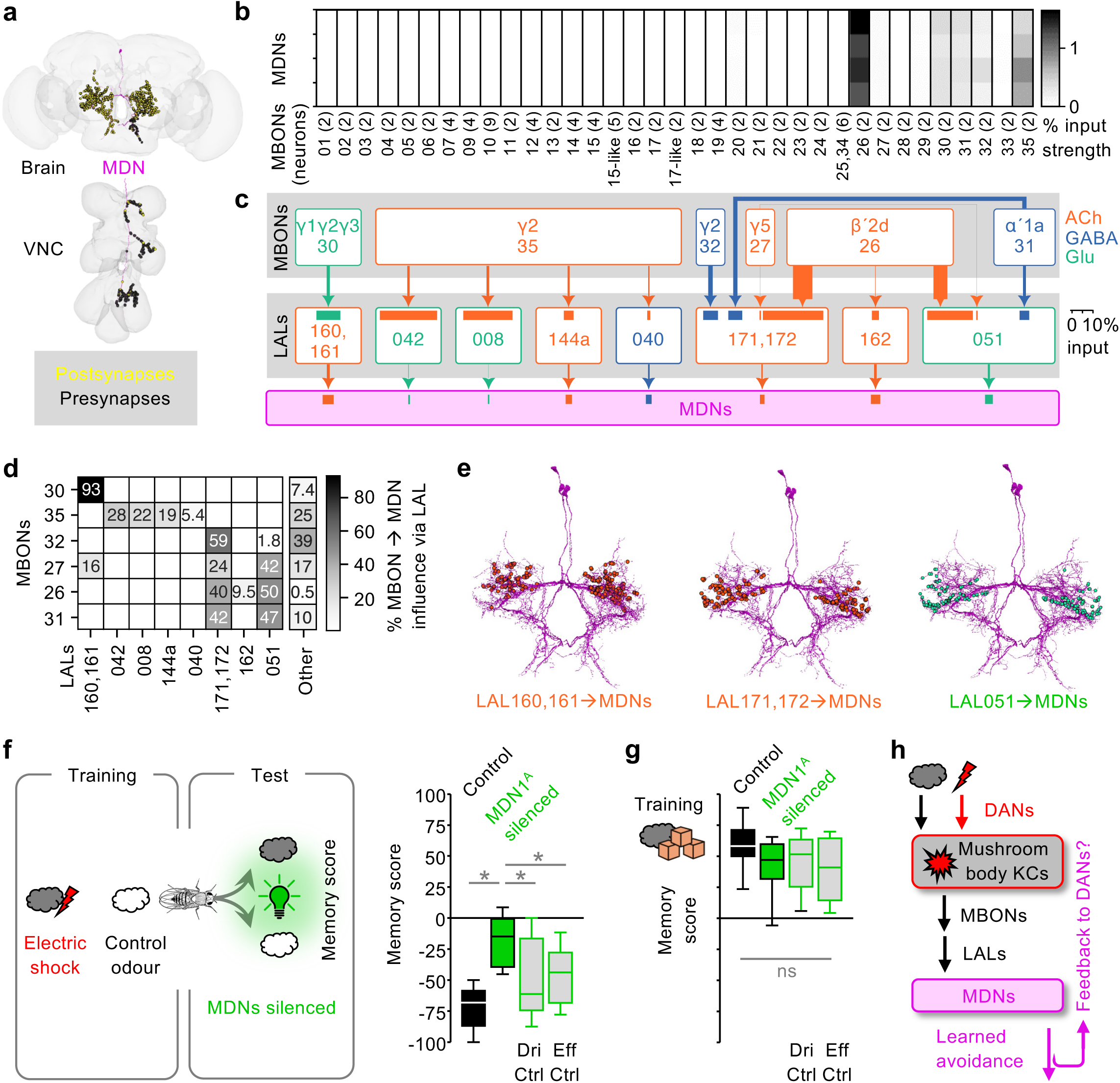
MDNs are part of aversive memory output pathways. **a,** EM reconstruction of a moonwalker descending neuron (MDN, magenta). VNC: ventral nerve cord. Yellow and black circles: post- and pre-synaptic sites, respectively. **b,** Heatmap of percent input for 2-step pathways reaching each MDN (rows) from each mushroom body output neuron type (MBONs, columns), determined from FlyWire-v783 after removing connections <10 synapses. Bracketed numbers refer to the number of neurons summed across hemispheres. **c,** Pathways from MBONs via neurons of the lateral accessory lobe (LALs) to the MDNs, combined for both hemispheres (omitting connections with <20 synapses total). Arrows are proportional to the total number of synapses (thinnest: 43 synapses, thickest: 2002 synapses). Horizontal bars show percent of input to the downstream partner, calculated as an average over the downstream neuron type, summed over the upstream neuron type. ACh: acetylcholine, orange. GABA: γ-aminobutyric acid, blue. Glu: glutamate, green. **d,** Percent of 2-step input to MDNs from each MBON type that passes through the indicated LAL. Percentages are calculated after averaging across MDNs. “Other” represents all other 2-step MBON-to-MDN pathways. **e,** Locations of synapses (dots) from the indicated LALs to the MDNs (magenta) colour-coded for transmitter as in **(c)**. **f,** Overview and outcome of odour-shock learning experiments. Clouds: odours. Lightning bolt: electric shock. Light bulb: optogenetic silencing of MDNs. Relative to the Control condition with MDN signalling intact (black), silencing MDNs during the test reduced odour-shock memory scores (MDN1^A^ silenced, green) (genotype in both cases: MDN1^A^>GtACR1) to levels less than in genetic controls (grey: Driver control: MDN1^A^>+, Effector control: +>GtACR1) (N= 13,22,21,21). **g,** As in **(f)**, but for pairings of odours with sugar reward (orange cubes), showing that appetitive memory scores remained unaffected (N= 8,9,8,10). **h,** Role of MDNs in aversive memory output pathways and hypothesis of feedback to DANs. Box-whisker plots show median, interquartile range (box) and 10th/90th percentiles (whiskers). Data were analysed across groups by Kruskal-Wallis tests (P< 0.05) (ns: P> 0.05), followed by pairwise comparisons (Mann-Whitney U-tests, *P< 0.05 with Bonferroni-Holm correction). Higher-resolution versions of **(a,e)** and additional information in Supplemental Data Synapse Topology. NeuronIDs for **(a-e)** in Methods Supplemental Table 2. Additional information in Extended Data Fig. 3.

**Fig. 3.**
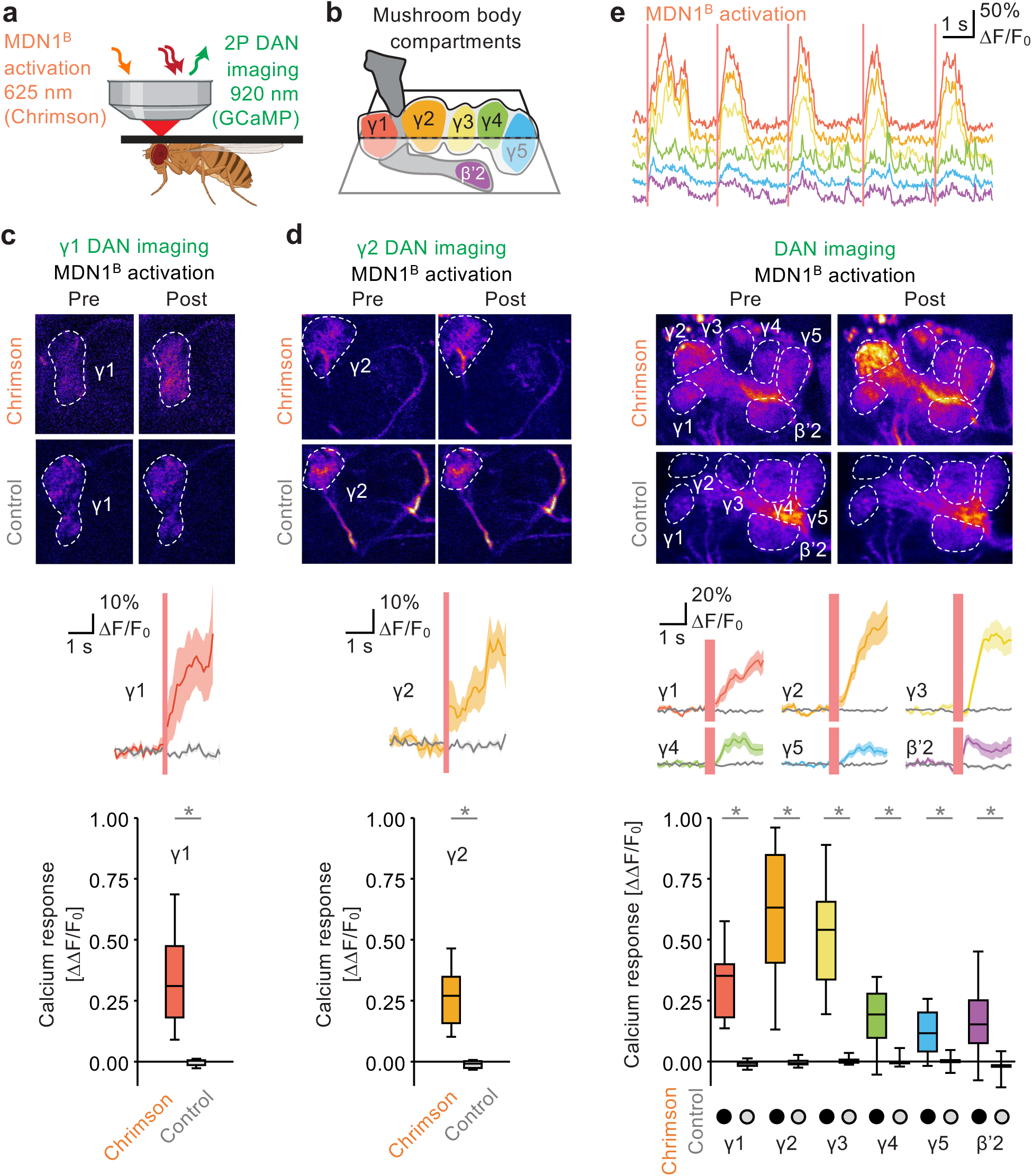
Activation of MDNs favours activity in punishing DANs. **a,b,** Combined optogenetics and *in vivo* imaging setup **(a)**. Red light pulses of 200 ms were used to activate Chrimson-expressing MDNs, while calcium signals were measured in GCaMP6f-expressing mushroom body DANs (**b**; compartments colour-coded) using continuous two-photon excitation scanning. **c,d,** Average intensity projections of sample recordings 2 s before (Pre) and 2 s after the first MDN activation (Post) in flies expressing Chrimson in γ1 **(c)** or γ2 **(d)** DANs, and in Control flies not expressing Chrimson (grey); dashed lines indicate compartment boundaries (top panels). Calcium transients (ΔF/F_0_) upon optogenetic MDN activation (red vertical bars) in flies expressing Chrimson (coloured traces) and in Controls (grey traces) (middle panels). Activation of MDNs results in significant calcium responses (ΔΔF/F_0_) in DANs of the γ1 (**c**; N= 6 flies each) and γ2 compartments (**d**; N= 8,7 in Chrimson and Control flies) compared to those in Controls (bottom panels). Experimental genotypes: MDN1^B^>Chrimson^C^; γ1>GCaMP **(c)** and MDN1^B^>Chrimson^C^; γ2>GCaMP **(d)**. Control genotypes: MDN1^B^>+; γ1>GCaMP **(c)** and MDN1^B^>+; γ2>GCaMP **(d)**. **e**, Sample traces of raw calcium transients (ΔF/F_0_) across the DANs of the compartments colour-coded as in **(b)** upon MDN activation (red bars) (top panel). Panels below as in **(c,d)** but expressing GCaMP across the DANs, revealing strong responses in γ1− γ3 (N= 10,8 in Chrimson and Control flies). Experimental genotype: MDN1^B^>Chrimson^C^; DANs>GCaMP. Control genotype: MDN1^B^>+; DANs>GCaMP. Box-whisker plots show median, interquartile range (box) and 10th/90th percentiles (whiskers). Except in the top panel in **(e)**, calcium transients are plotted as mean ± SEM. Data were analysed by Mann-Whitney U-tests (*P< 0.05) **(c,d)** or by Kruskal-Wallis tests (P< 0.05), followed by pairwise comparisons (Mann-Whitney U-tests, *P< 0.05 with Bonferroni-Holm correction) **(e)**. Quantification in the bottom panels in **(c-e)** is based on the first optogenetic activation. Additional information in Extended Data Fig. 4,5 and Supplemental Video 1,2.

### MDNs are part of aversive memory output pathways

MDN dendrites innervate the lateral accessory lobe (LAL), where most of their inputs are received, whereas most of their axonal output terminals are in the sub-oesophageal zone and the ventral nerve cord (Fig. 2a; Supplemental Data Synapse Topology). Our analysis of the fly chemical synapse connectome (Scheffer et al. 2020, Dorkenwald et al. 2024, Schlegel 2024) suggests that the MDNs are mainly influenced by only 6 of the 35 types of mushroom body output neuron (MBONs) (Fig. 2b,c), all of which are classified as atypical (Li et al. 2020). The strongest influence is exerted by MBON26 and MBON35 (Fig. 2b) and is mediated by 8 types of local neuron in the LAL, with LAL160,161 as well as LAL171,172 and LAL051 as the main hubs (Fig. 2c-e; Supplemental Data Synapse Topology). A substantial share of the synaptic pathways from the mushroom body to the MDNs involve MBONs of the punishment-memory compartments γ1-γ3 (MBON30, MBON35, MBON32). Moreover, 4-5 of these 6 MBONs, as well as LAL171,172 and LAL051, have previously been suggested as circuit elements by which associative memories are behaviourally expressed (MBON26, MBON27, MBON31, MBON32 and likely MBON35: Li et al. 2020). We therefore tested whether the MDNs are part of memory-efferent pathways.

Flies underwent differential conditioning with odours as conditioned stimuli and electric shock as the unconditioned stimulus, an established Pavlovian association paradigm that produces aversive olfactory short-term memory local to the punishment compartments of the mushroom body (Heisenberg 2003, Gerber & Aso 2017, Cognigni et al. 2018, Boto et al. 2020, Modi et al. 2020, Davis 2023). When MDNs were optogenetically silenced during the choice test, the behavioural expression of odour-shock memory was massively impaired, whereas no such effect was seen in the genetic controls (Fig. 2f). Notably, the behavioural expression of appetitive odour-sugar memory was not likewise affected (Fig. 2g), and neither was innate olfactory choice behaviour (Extended Data Fig. 3). These results show that the MDNs contribute to the behavioural expression of aversive odour-shock memory (Fig. 2h).

We next focused on how, in turn, MDNs affect mushroom body processing. Given that activation of MDNs in the presence of an odour produces an aversive memory for that odour (Fig. 1j), we asked whether the activation of MDNs drives feedback to punishing dopaminergic neurons (DANs) (Fig. 2h).

### Activation of MDNs favours activity in punishing DANs

To test for MDN-to-DAN feedback, we combined optogenetic activation of MDNs with *in vivo* two-photon calcium imaging of DANs in the mushroom bodies’ γ1 and γ2 compartments, known to mediate punishing effects (Claridge-Chang et al. 2009, Aso et al. 2010, Aso et al. 2012). We generated transgenic flies that allowed us both to optogenetically activate MDNs via the red-light-activated channelrhodopsin Chrimson and simultaneously to image specific DANs by the calcium indicator GCaMP6f through a small window cut into the dorsal head capsule. After mounting the flies under a two-photon microscope with their heads, thorax and wings fixed but their legs free to move (Fig. 3a,b), we observed that activation of MDNs resulted in strong calcium responses in both the γ1 DAN (PPL1-01: Fig. 3c) and the γ2 DAN (PPL1-03: Fig. 3d) (Extended Data Fig. 4). Neither in these nor in any of the following experiments did we observe calcium responses in control flies without the optogenetic effector.

To confirm these observations regarding the γ1 and the γ2 DAN, and to extend our focus beyond these two ‘usual suspects’, we used a fly strain that expresses the GCaMP6f calcium reporter more broadly across the DANs and thus permits signals to be monitored in all five compartments of the γ lobe and the β’2 compartment (Fig. 3b). Upon activation of MDNs, we saw the strongest responses in the DANs of the γ1, γ2, and lateral regions of the γ3 compartment, all of which have previously been shown to mediate punishing effects (Claridge-Chang et al. 2009, Aso et al. 2010, Aso et al. 2012, Li et al. 2020), whereas responses in rewarding DANs (medial γ3, γ4, γ5, and β’2 compartments: Burke et al. 2012, Liu et al. 2012, Li et al. 2020) were considerably weaker (Fig. 3e; Extended Data Fig. 5; Supplemental Video 1, 2).

These results show that activation of MDNs favours activity in punishing over rewarding DANs, suggesting a mechanism by which MDN activation produces aversive memories for concomitantly presented odours. We next asked whether such DAN activation is mediated by internal, recurrent feedback from MDNs to DANs, or whether the execution of MDN-evoked movement generates external sensory feedback that in turn activates DANs.

### No evidence for internal, recurrent feedback from MDNs to DANs

We initially hypothesized that there is internal, recurrent feedback from MDNs to DANs, but systematic queries of the brain connectome of the fly (Scheffer et al. 2020, Dorkenwald et al. 2024, Schlegel 2024) did not reveal any credible such pathway (see Methods section for inclusion criteria). Current knowledge of ascending input from the ventral nerve cord to the brain does not offer evidence for a pathway from the MDNs to the DANs either (Azevedo et al. 2024, Stürner et al. 2024). We therefore tested whether an as yet unidentified internal MDN-to-DAN pathway might be functionally relevant. When we optogenetically activated MDNs and imaged the DANs in an explant brain or in a brain-plus-ventral-nerve-cord preparation, no activation of any DAN was observed, however (Extended Data Fig. 6). We therefore considered external, reafferent feedback from the execution of MDN-evoked movement.

### Movement is required for DAN activation and punishment by MDNs

We returned to a combination of optogenetic activation of MDNs with *in vivo* calcium imaging of DANs but this time transiently restraining leg movements (Fig. 4). Under conditions of unrestrained leg movements, we confirmed reliable and strong responses of DANs in the γ1-γ3 compartments (Before trapping; Fig. 4a,d,e). These responses were abolished when leg movements were transiently restrained by a piece of cotton wool gently applied to the fly (Trapped; Fig. 4b,d,e), and largely recovered after removing the restraint (After trapping; Fig. 4c-e) (Supplemental Video 3). These results show that the execution of MDN-evoked leg movements is required for activation of the DANs of the γ1-γ3 compartments. Indeed, the onset of leg movements precedes their activation by as much as 500 ms (Extended Data Fig. 7), consistent with reafferent, sensory feedback from executed leg movements being the cause of this activity, but longer than plausible for internal, recurrent MDN-to-DAN feedback.

**Fig. 4.**
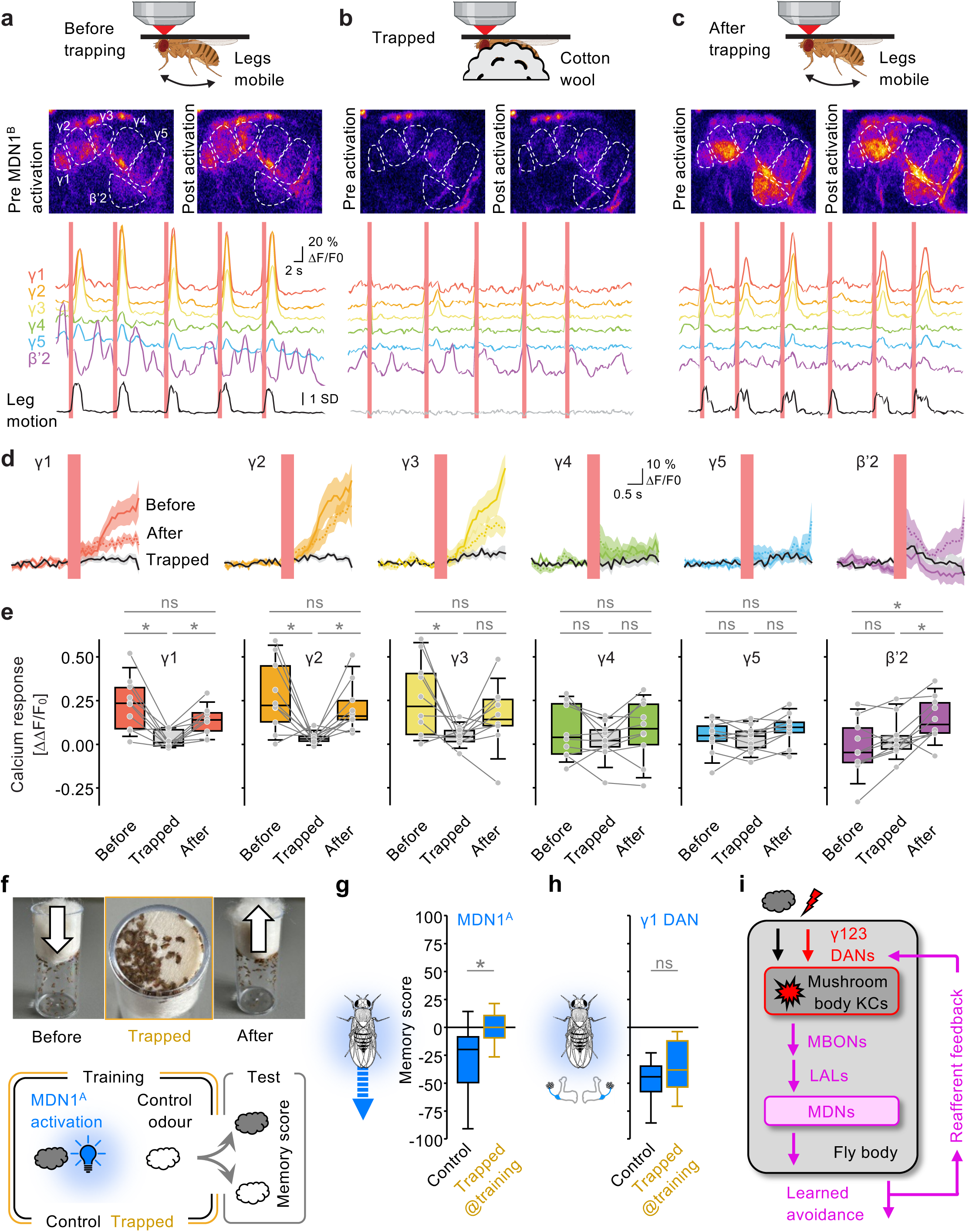
Movement is required for DAN activation and punishment by MDNs. **a-c**, Two-photon *in vivo* imaging before **(a)**, while **(b)**, and after **(c)** leg movement was restrained (Trapped) using a piece of cotton wool (top panels). 200 ms of red light stimulation was used to activate MDNs via Chrimson, while calcium signals were measured in the DANs of the indicated mushroom body compartments with GCaMP6f. Leg movements were calculated as the legs’ motion energy in videos captured by an infrared camera. Average intensity projections of the same field of view across conditions, from 2 s before (Pre) and 2 s after MDN activation (Post); dashed lines indicate compartment boundaries (middle panels). Sample traces of raw calcium transients (ΔF/F_0_) in the DANs of the indicated compartments and leg motion energy (bottom panels) with MDN activation indicated by red vertical bars. **d**, Calcium transients (mean ± SEM) in DANs of the indicated compartments upon the first MDN activation (red vertical bars) before (coloured lines), while (black lines, Trapped), and after (dotted lines) leg movement was restrained. **e**, MDN-evoked calcium responses (ΔΔF/F_0_) in DANs of the indicated compartments recorded across trapping conditions. Data points from individual flies are connected by lines (N= 10 flies). Genotype **(a-e)**: MDN1^B^>Chrimson^C^; DANs>GCaMP. **f**, Procedure for transiently restraining movement in the behavioural setup (top, Trapped) and overview of learning experiments (bottom). Clouds: odours. Light bulb: optogenetic activation of neurons indicated in **(g, h)**. **g,h**, Pairing odour with optogenetic activation of MDNs establishes aversive memory under Control conditions but not when movement was restrained during the training period (**g**). No effect of such restraint was observed when activating the DANs of the γ1 compartment **(h)**. Genotypes: MDN1^A^>ChR2XXL^A^ (N= 29,29) **(g)** and γ1>ChR2XXL^A^ (N= 24,24) **(h)**. **i**, Schematic of reafferent feedback from learned avoidance to dopaminergic teaching signals. Box-whisker plots show median, interquartile range (box) and 10th/90th percentiles (whiskers). Data were analysed by Wilcoxon signed-rank tests with Bonferroni-Holm correction **(e)** or Mann-Whitney U-tests **(g, h)** (*P< 0.05) (ns: P> 0.05). Additional information in Extended Data Fig. 6-8 and Supplemental Video 3,4.

We next asked whether the execution of MDN-induced movement is also required for the punishing effect of MDN activation, using a procedure to transiently restrain movement during training (Fig. 4f) (Supplemental Video 4). Flies were trained differentially such that one but not the other odour was paired with optogenetic activation of the MDNs, followed by a choice test between the two odours. This confirmed the punishing effect of activation of the MDNs (Fig. 1j; Extended Data Fig. 4) – but only when the training took place under control conditions such that the flies could freely move in the training apparatus (Control; Fig. 4g). In contrast, no memories were formed when the flies were gently restrained in their movement during the training period (Trapped@training; Fig. 4g). No adverse effect of restraint was observed when known punishment DANs were activated directly (Fig. 4h), showing that in principle aversive olfactory memory formation is possible under these conditions. To our initial surprise, but consistent with the lack of spontaneous DAN activity under restraint (Fig. 4b), restraint is not punishing in itself (Extended Data Fig. 8a).

These results show that the execution of MDN-evoked movements is required both for MDN activation to engage the punishing γ1-γ3 DANs, and for MDN activation to have a punishing effect. Together with our earlier results, this demonstrates reafferent feedback from learned avoidance to the teaching signals establishing aversive memory (Fig. 4i) – raising the question of the adaptive significance of such feedback.

### MDN-mediated feedback counterbalances extinction learning

We hypothesized that feedback from learned avoidance to aversive teaching signals counter-balances extinction learning. That is, after pairing a conditioned stimulus (CS) and a punishing unconditioned stimulus (US), animals typically retreat from the CS to avoid the US. Such avoidance breaks the initially learned CS-US contingency and should therefore initiate extinction learning. However, despite the broken CS-US contingency, learned avoidance of the CS is often maintained across multiple encounters, a phenomenon referred to as the “avoidance paradox” (Bolles 1972, Le Doux et al. 2017). We hypothesized that MDN feedback facilitates this maintenance of avoidance and developed a reinforcement learning model to probe its performance with or without MDN feedback, as well as an experimental test of this notion.

In the model, connections from odour-responsive KCs onto two representative MBONs that promote approach or avoidance are modulated by two representative DANs that are activated by punishment and reward, respectively (Fig. 5a). DANs receive signals encoding external reinforcement as well as predicted reinforcement calculated from MBON activity. Together, these DANs represent the prediction error of standard reinforcement learning models. In addition, in our model the DANs receive reafferent feedback from MDN-mediated learned avoidance, that is, when the learned value of the odour is negative and avoidance behaviour is executed.

**Fig. 5.**
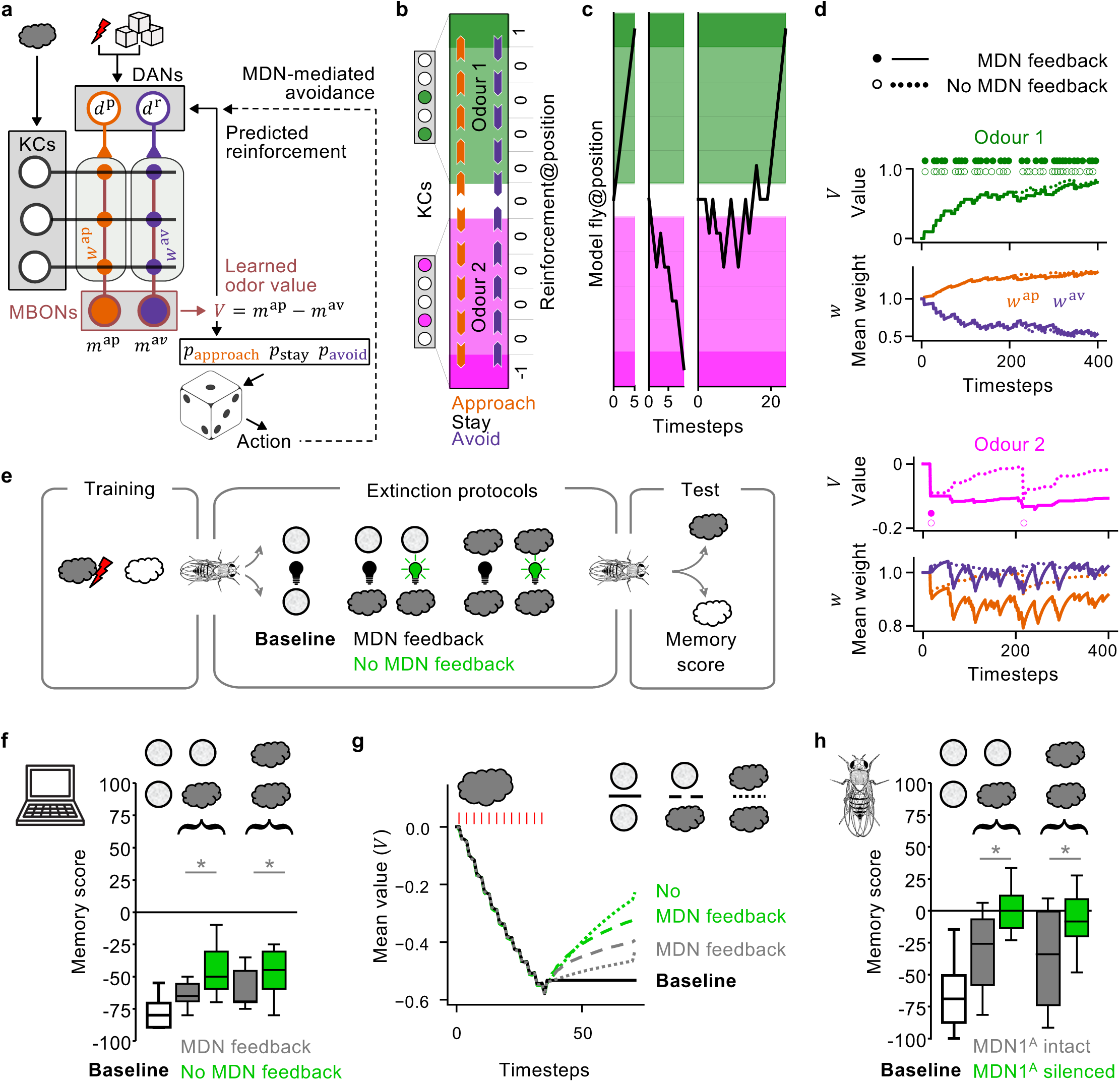
MDN-mediated feedback counterbalances extinction learning. **a,** Model schematic. Punishment and reward activate DANs with firing rates *d*^p^ and *d*^r^, respectively. MBONs with firing rates *m*^ap^ and *m*^av^promote approach and avoidance, respectively. KC-to-MBON weights (*w*^ap^, *w*^av^) are depressed by co-activation of DANs and odour-responsive KCs. Approach or avoidance behaviour is determined probabilistically by the value (*V*) derived from MBON activity. Dice by Steaphan Greene, CC-BY-SA-3.0. **b,** Schematic of one-dimensional model environment. Reinforcement (punishment -1, reward 1) is received at either end. Different odours (green, magenta) activate different KCs. Arrowheads show possible action choices. **c,** Example trials of a single model fly navigating the arena. Trials end upon first reinforcement (left and right: reward, middle: punishment). Left and middle are before learning, right is after learning. **d,** Evolution of model parameters over multiple trials of the paradigm shown in **(b)**. Circles denote ends of trials when the model fly received reward (top, odour 1) or punishment (bottom, odour 2). The value (*V*) of the rewarded odour 1 is learned through changes in KC-to-MBON weights (*w*^ap^ and *w*^av^, averaged across odour-responsive KCs) regardless of MDN feedback. Without MDN feedback, avoidance of the punished odour 2 is less persistent and the model fly receives a second punishment earlier (∼timestep 200). **e,** Schematic of training, extinction protocols and test in the five experimental conditions. Clouds: odours. Grey circle: choice option without odour. Black and green light bulbs: conditions with or without MDN feedback. **f,** Results of simulations of protocols in **(e)**. Without MDN feedback the full extent of extinction learning is revealed. Results averaged over 20 experiments per protocol, with 20 model flies per experiment. **g,** Mean learned odour value (*V*) during the protocols in **(e, f)**. During acquisition (timesteps 1-36), 12 pulses of punishment (red bars) are delivered in the presence of odour. During the extinction protocol (after timestep 36), MDN feedback counterbalances the return of odour value to pre-training levels. Line thickness exceeds ±2 s.e. **h,** Behavioural experiment as in **(e, f)**, using optogenetic silencing of the MDNs. Genotype: MDN1^A^>GtACR1 (N= 50,34,31,18,16). Box-whisker plots show median, interquartile range (box) and 10th/90th percentiles (whiskers). Data were analysed by Kruskal-Wallis tests (P< 0.05), followed by pairwise comparisons (Mann-Whitney U-tests, *P< 0.05 with Bonferroni-Holm correction). Additional information in Extended Data Fig. 8,9.

We tested model flies in a one-dimensional virtual environment with two odours (Fig. 5b,c). Reward or punishment is paired with either of these odours and is received only when the model fly reaches the edge of the environment. At each time-point, the model fly chooses either to stay in its current location, to approach, or to avoid the odour it senses. In the case of odour-reward pairings, the behaviour of model flies is not affected by the presence or absence of MDN feedback (Fig. 5d, top). Disabling MDN feedback has drastic consequences in the aversive domain, however. Without MDN feedback and thus with extinction learning operating as in conventional reinforcement learning models, updates to KC-MBON connection weights, odour value and learned avoidance quickly return to pretraining levels (Fig. 5d, bottom). As a result, the model fly soon receives additional punishment. By contrast, in model flies with MDN feedback intact, dopamine transients that reinforce the negative value of the punished odour each time an avoidance action is taken maintain the avoidance memory and prevent the flies from encountering future punishment (Fig. 5d, bottom).

The above modelling results suggest that MDN feedback counterbalances extinction learning. We therefore simulated model flies undergoing aversive conditioning followed by extinction protocols, either with MDN feedback intact or without it (Fig. 5e). After an extinction protocol with MDN feedback intact, intermediate memory scores were observed. Without such feedback, memory scores were further reduced, revealing the full effects of extinction learning (Fig. 5f,g). These model results hold both in situations when, during the extinction protocol, an opportunity is given to avoid the punishment-predicting odour, and when the model flies can only choose between two equally punishment-predicting options (the latter protocol eliminates differences in odour exposure with or without MDN feedback).

We experimentally tested these model predictions, using corresponding behavioural paradigms in flies with MDNs intact or optogenetically silenced during the extinction protocol. Memory scores in these behavioural experiments qualitatively matched those predicted by the model (Fig. 5h; silencing the MDNs does not in itself confer valence: Extended Data Fig. 8b).

Together, the model and the behavioural experiments demonstrate that an adaptive significance of reafferent feedback from learned avoidance to punishing teaching signals is that it counterbalances extinction learning and maintains successful learned avoidance.

## Discussion

### Error types and the avoidance paradox

Inspired by the hypothesis of mutual causation between action and valence (Darwin 1872, James 1884), we discovered that the execution of avoidance manoeuvres engages punishing dopaminergic teaching signals that can induce aversive memories even if no external punishment is received (Fig. 1-4; Extended Data Fig. 4,5). This process enables the continuation of avoidance to save the animal from receiving another, potentially lethal, punishment (Fig. 5; Extended Data Fig. 9a). However, continued avoidance in a situation that is benign would be maladaptive. The two types of error that can result from these policies, i.e. no longer avoiding even though avoidance is warranted or avoiding unnecessarily, cannot both be simultaneously minimized, as is reflected in the “avoidance paradox” (Bolles 1972). We suggest that two separate systems control the trade-off between these errors. In the γ1-γ3 compartments, learned avoidance engages aversive teaching signals to maintain aversive odour memory and promote future avoidance (this study), thus reducing the former error. In the γ5 compartment, extinction memories that oppose further avoidance of the odour are established (Felsenberg et al. 2018), reducing the latter error. This allows these policies to be selected according to situational, motivational, and mnemonic variables.

Our study thus reveals that, during extinction protocols, aversive memories are not merely unaffected or left to decay but instead are actively maintained. Enacted avoidance engages punishing teaching signals that sustain memory for the cues that had triggered the avoidance. This qualitatively extends the notion of the parallel neuronal organization of initially acquired aversive memory and memories established by extinction learning (Tovote et al. 2015, Felsenberg et al. 2018) (Extended Data Fig. 9b,c). It is furthermore compatible with the proposal that successful avoidance establishes a state of relief that reinforces the avoidance behaviour as the action that brought about the relief (Lloyd & Dayan 2026, LeDoux et al. 2017, Bouton et al. 2021, Laing et al. 2025) (in flies, avoidance-promoting MBONs seem poised to receive such reinforcement). In concert, these learning processes can ensure that when the cue is encountered again, the action that prevented the punishment is repeated.

### Memory-efferent pathways

A selective set of MBONs and LALs with acetylcholine, GABA or glutamate as their predicted transmitters establishes 2-step connections from the MBONs to the MDNs (Fig. 2). We here assume that acetylcholine has the excitatory and GABA the inhibitory effects typical of insect central brain synapses; for glutamate, inhibitory effects are assumed (Shiu et al. 2024, based on Liu & Wilson 2013). Accordingly, all but one of the pathways originating in MBONs of the punishment compartments γ1-γ3 feature either an excitatory MBON upstream of an inhibitory LAL, or vice versa (Fig. 2c). Through such sign-inversion, a learning-induced depression of KC-MBON synapses will promote MDN activity and backward movement as an early component of avoidance before the fly turns around and assumes a new direction (Extended Data Fig. 10).

Silencing the MDNs during the choice test impaired but did not abolish the behavioural expression of odour-shock memory (Fig. 2f). This suggests that learned avoidance can be expressed by pathways parallel to the MBON-LAL-MDN pathways (Fig. 2b,c). This has indeed been shown for the MBON of the γ1 compartment (MBON-11) and its target MBONs of the γ5 and β’2 compartments (MBON-01, MBON-03) (Aso et al. 2014, Owald et al. 2015, Perisse et al. 2016). Such a parallel organization highlights the complexity and importance of the decision whether to avoid.

### Reafferent pathways and DAN signalling

We demonstrated reafferent feedback from MDN-induced movement to aversive dopaminergic teaching signals (Fig. 4). Which sensory modality registers these movements? Visual input from movement-over-ground is likely to be irrelevant because functional imaging experiments did not allow for such movement. More probable is a role for leg proprioceptive organs (campaniform sensilla, hair plates, chordotonal organs: Tuthill & Azim 2018, Büschges & Ache, in press). Indeed, optogenetic activation of the chordotonal organs selectively activates punishing DANs in larval *Drosophila* (Eschbach et al. 2020).

The feedback from MDN-induced movement to punishing DANs (Fig. 4,5) adds complexity to the picture of mushroom body DAN function (Riemensperger et al. 2005, Cohn et al. 2015, Hattori et al. 2017, Felsenberg et al. 2018, Li et al. 2020, Otto et al. 2020, Siju et al. 2020, Driscoll et al. 2021, Jiang & Litwin-Kumar 2021, Zolin et al. 2021, Cazalé-Debat et al. 2024, Meschi et al. 2024). As a heterogeneous population, the mushroom body DANs establish a nuanced, combinatorial coding space for salient features of the animal’s present and predicted environment, its current goals and action tendencies, as well as its situationally relevant past experiences. Collectively, these influences shape present and future action selection. Similar heterogeneity is observed in dopamine neurons in the ventral tegmental area of mammals (da Silva et al. 2018, Stuber 2023). Given this complexity and given that the circuit position of the mushroom body DANs is far removed from the sensory and motor periphery, the relationship between movement and DAN activity is heavily modulated by situational, behavioural and motivational variables (Siju et al. 2020, Zolin et al. 2021).

### Implications

Extinction is a central component of exposure therapies, serving as effective first-line treatments of, for example, anxiety disorders. However, resilience to post-therapy adverse experiences and generalization beyond the therapeutic context are not yet satisfactory (Vervliet et al. 2013, Carpenter et al. 2018, Craske et al. 2018). Correspondingly, in rodents, extinguished aversive behaviours often return upon re-exposure to punishment (reinstatement) or upon contextual change (renewal) (Bouton et al. 2021, Laing et al. 2025). The engagement of punishment signals through avoidance behaviour as reported in the present study seems poised to maintain aversive memories in a state susceptible to such reinstatement and renewal. Extrapolated to the human condition, the clinical implication is that preventing avoidance during exposure therapy can reduce relapse rates because it prevents the engagement of avoidance-induced punishment signalling. Conceptually, our findings call for an integrated view of the brain mechanisms of behaviour organization, memory function and emotion regulation.

## Supporting information

Supplemental Data Synapse Topology

## Methods

### Fly strains

*Drosophila melanogaster* were raised under standard conditions, in constant darkness to avoid any unintended optogenetic effects. For behavioural experiments, mixed-sex cohorts of 2-5-day-old flies were used. Calcium imaging and immunohistochemistry used female flies only due to genetic constraints. Strains to establish experimental genotypes and their driver and effector controls (Dri Ctrl and Eff Ctrl) have been described previously, unless otherwise mentioned. For further details, see Supplemental Methods Table 1.

### Conditioning and choice experiments

Male flies of the driver strains were crossed to females of the effector strains, and cohorts of ∼60-100 flies of their F1 progeny were used in a T-maze setup as described previously (Schwaerzel et al. 2002) (CON-ELEKTRONIK, Greussenheim, Germany), allowing for electric shock, optogenetic manipulation and odour delivery, operated at 23-25 °C, 60-80 % relative humidity, and red light invisible to flies, unless mentioned otherwise. As odorants, undiluted 50 µl benzaldehyde (BA) and 250 µl 3-octanol (OCT) (CAS 100-52-7, 589-98-0; Fluka, Steinheim, Germany) were applied to Teflon containers of 5 mm and 14 mm diameter, respectively.

#### Conditioning with optogenetic activation as reinforcement

Olfactory conditioning with optogenetic activation as reinforcement was conducted as described previously (König et al. 2018) (Fig. 1b). Flies were loaded into the training tubes and 2 min later one odour (CS+) was presented for 1 min. Unless otherwise mentioned, 15 s later (inter-stimulus-interval (ISI) of -15 s) pulsed light for optogenetic activation was turned on for 1 min (training tubes featured either 24 blue LEDs (465±10 nm, 0.069 mW/mm^2^) or 24 red LEDs (627±10 nm, 0.053mW/mm^2^)) at 4.99 s / 0.01 s ON / OFF pulses. Another 4 min later the control odour (CS-) was presented. In total, flies underwent three such training trials. Then they were given a choice test between the two odours loaded to the arms of the T-maze. After 2 min, the arms of the maze were closed, and relative odour preference was calculated from the number of flies (#) in each arm:

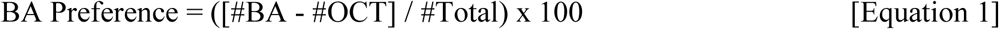

Across repetitions of the experiment, flies were trained with contingencies of odour and light (*) swapped to calculate the memory score:

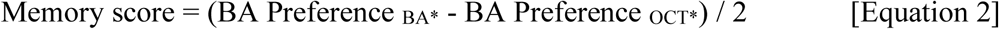

Positive memory scores thus indicate appetitive and negative memory scores aversive associative memory. The experiment in Extended Data Fig. 8b was performed the same way, except that green light (530±10 nm, 0.037mW/mm^2^) was used for optogenetic silencing.

#### Behavioural pharmacology

Pharmacological manipulations were performed as described previously (Amin et al. 2025). For 36-40 h before behavioural experiments flies were offered as their sole food a tissue paper (Fripa, Düren, Germany) soaked with 1.8 ml of either (i) a plain 5 % sucrose solution (CAS: 57-50-1, Hartenstein, Würzburg, Germany), or that solution added with either (ii) 5 mg/ml 3-iodo-L-tyrosine (3IY), an inhibitor of dopamine synthesis (CAS: 70-78-0, Sigma, Steinheim, Germany), or (iii) 5mg/ml 3IY plus the dopamine precursor 10 mg/ml 3,4-dihydroxy-L-phenylalanine (L-DOPA) (CAS: 59-92-7, Sigma, Steinheim, Germany). In all cases, flies were trained and tested as described above for optogenetic activation as reinforcement, at ISIs of either -15 s or 120 s (Fig. 1h).

#### Conditioning with shock or sugar reinforcement and optogenetic silencing during the test

Olfactory conditioning with electric shock or sugar as reinforcement was conducted as described above for optogenetic activation as reinforcement with the following differences. Flies went through only one training trial either with electric shock (12 pulses of 100 V, 1.2 s / 3.8 s ON / OFF) or with 2 M sucrose solution presented on filter paper. During the 2 min of the choice test, green light was turned on for optogenetic silencing (Fig. 2f,g). Innate olfactory choice behaviour was assessed the same way, save for the training (Extended Data Fig. 3).

#### Restraining movement (trapping) during training

For the Control case, olfactory training and testing was performed with optogenetic activation as reinforcement as described above. For the Trapped case, flies were gently pushed down to the bottom of the training tubes using a piece of cotton wool, and released from this restraint before the choice test (Fig. 4f,g, Supplemental Video 4). The experiment in Fig. 4h used only one training trial to avoid overtraining (König et al. 2018). The experiment in Extended Data Fig. 8a used one training trial of olfactory conditioning with trapping as reinforcer.

#### Extinction learning

Extinction learning experiments were modified from a previously published procedure (Felsenberg et al. 2018, Schwaerzel et al. 2002) (Fig. 5e). Training with odour and electric shock was conducted as described above. Subsequently, flies were removed from the T-maze setup and kept on a standard food vial for 30 min before being returned to the setup and released at the choice point. For the flies of the baseline condition, only air was presented on both sides of the T-maze for 1 min, whereas independent sets of flies underwent extinction protocols such that the CS+ was presented either in one arm or in both arms of the T-maze. These extinction protocols were run, in independent sets of flies, without green light or with pulsed green light for optogenetic silencing as mentioned before (MDN feedback, and No MDN feedback, respectively). After being collected from both arms of the maze, flies were subjected to a 2-min choice test between CS+ and CS-as described above.

### *In vivo* two-photon calcium imaging

A custom-made two-photon microscope with a protocol modified from Barnstedt et al. (2016) was used. 2-7-day-old flies of the indicated genotypes were transferred to freshly prepared standard food vials mixed with 1mM all-trans-retinal for 2-3 days. Flies were briefly (∼20 s) anaesthetized on ice and tethered in a custom chamber to open the head capsule under room temperature in carbogenated (95 % O_2_, 5 % CO_2_) buffer solution (103 mM NaCl, 3 mM KCl, 5 mM N-Tris, 10 mM trehalose, 10 mM glucose, 7 mM sucrose, 26 mM NaHCO_3_, 1 mM NaH2PO_4_, 1.5 mM CaCl_2_, 4 mM MgCl_2_, osmolarity 275 mOsm, pH 7.3-4). Flies were fixated using wax attached to eyes, wings and parts of the thoracic body wall, but were able to move their legs. After we had confirmed genetic markers and transgene expression under a tabletop fluorescence microscope (Extended Data Fig. 5d), flies were transferred to the two-photon setup.

For optogenetic activation, a high-power red light 700mW LED (M625L4, Thorlabs) of 625±10 nm using the LEDD1B T-Cube LED Driver (Thorlabs) was relayed through the imaging objective onto the specimen, triggered manually. The power at the specimen was measured to be 10.5 mW/mm^2^. After preparation and transfer to the two-photon microscope, flies were left to rest for 3-5 min. 10 s after recording baseline fluorescence, five 200 ms (Fig. 3c-e, Fig. 4a-e, Extended Data Fig. 5b-b’’) or 20 ms (Extended Data Fig. 5c-g’’) light pulses were delivered at 40 Hz while recording calcium signals for a total of 60 s. Fluorescence was excited using 75 fs pulses, 80 MHz repetition rate, centred on 920 nm generated by a Ti-Sapphire laser (Chameleon Vision S, Coherent), and 550 x 550 pixel images were acquired at 40 Hz, controlled by custom-written software in LabView (National Instruments). Dopaminergic neurons were imaged at the level of the mushroom body horizontal lobe, focusing on the γ1− γ5 and β’2 compartments. Regions of interest (ROI) were manually drawn using ImageJ and analysed using Microsoft Excel, Statistica and custom-written Python scripts. Baseline fluorescence F_0_ was defined as the mean F from the first 2 s of recording. Calcium responses were quantified by comparing the average ΔF/F_0_ from 2 s before until light onset (pre) and the average ΔF/F_0_ from light offset to 2 s thereafter (post). ΔΔF/F_0_ was defined as the post-stimulus average fluorescence minus the pre-stimulus average fluorescence. For Extended Data Fig. 5, data from the γ3 subcompartments were similarly extracted from those brains with suitable orientation in the x axis (medial-lateral) to allow within-compartmental analyses, averaged across the y axis (anterior-posterior) and separated into 5 bins across the x axis.

For the experiment in Fig. 4a-e, leg movements were monitored using a monochrome CCD camera (Basler acA 780-75gm) positioned approximately 5-10 cm from the fly. Camera position was aligned for each fly before the start of recordings, and images were acquired at 75 Hz with 520 x 520 pixels using Pylon Camera Software Suite (Basler). Videos were synchronized to two-photon imaging recordings by the two-photon laser’s visible on- and offset. Files were saved in compressed MP4 format before further processing.

### Explant two-photon calcium imaging

Flies of the indicated genotypes were anaesthetized on ice for 5 mins and dissected in carbogenated buffer solution (103 mM NaCl, 3 mM KCl, 5 mM N-Tris, 10 mM trehalose, 10 mM glucose, 7 mM sucrose, 26 mM NaHCO_3_, 1 mM NaH2PO_4_, 1.5 mM CaCl_2_, 4 mM MgCl_2_, 295 mOsm, pH 7.3). Either only the brain or the brain with the ventral nerve cord attached was extracted (Extended Data Fig. 6a,b). Samples were put into a transparent imaging chamber and kept in place by a nylon grid. The chamber was then filled with buffer solution. After 5 min, recordings started under a Femtonics two-photon microscope using 920 nm pulsed laser light with a 30 Hz imaging rate. Optogenetic activation was performed with a Thorlabs red light LED of 625 nm (0.014 mW/mm^2^) controlled by a HEKA patch master EPC9 v2×91. Due to the orientation of the samples, the γ1 and β’2 compartments are located in a different focal plane from the γ2− γ5 compartments. Therefore, using three 200-ms pulses of red light for optogenetic activation, two z-planes were recorded starting either with γ1 and β’2, or the γ2− γ5 plane. Starting with the respective second plane, we repeated this for 1000-ms pulses of red light. Raw imaging files were downsampled to 15 Hz and converted into TIFF files using a custom-written Python script. Using NOSA (Oltmanns et al. 2020), changes in the fluorescence signal of the respective compartments were determined and exported as Excel files. Data were further analysed and visualized using custom-written Python scripts. Baseline fluorescence F_0_ was defined as the mean F from the first 2 s of recording. Calcium responses of the first stimulation trial were quantified by comparing the average ΔF/F_0_ from 2 s before until light onset (pre) with the average ΔF/F_0_ from light offset to 2 s (Δ long) or 0.5 s (Δ short) thereafter (post). ΔΔF/F_0_ was defined as the Δ short or Δ long post-stimulus average fluorescence minus the pre-stimulus average fluorescence.

### Immunohistochemistry

Moonwalker^A^ and MDN1^A^ were crossed to Chrimson^A^ (Fig. 1a,j, Extended Data Fig. 1a,c), whereas Moonwalker^B^ and MDN1^B^ were crossed to Chrimson^D^ (Extended Data Fig. 4a,b). Brains and ventral nerve cords from adult F1 progeny (5-7 days old) were dissected and immunostained as described (Wu et al. 2016) (https://www.janelia.org/project-team/flylight/protocols). Tissues were dissected in PBS on ice, fixed in 4 % PFA (20-30 min at room temperature), and washed in 0.5 % PBST (3 x 10-15 min at room temperature). After being left overnight in blocking solution (10 % NGS in PBST) at 4 °C, tissues were incubated with the primary antibody for 24 h at 4 °C and washed in 0.5 % PBST (3 x 10-15 min at room temperature, and overnight at 4 °C). Tissues were incubated with the secondary antibody for 24 h at 4 °C and washed in 0.5 % PBST (3 x 10-15 min at room temperature, and overnight at 4 °C). After a final washing step (1 x 5-10 min in PBS), tissues were mounted on a slide using the DPX mounting protocol (https://www.janelia.org/project-team/flylight/protocols). Image z stacks were acquired with a LSM780 confocal microscope (Zeiss NY, USA) at 1024 × 1024 pixel resolution. Image processing was performed using ImageJ (version 1.53f, Fiji ImageJ).

For Moonwalker^A^ and MDN1^A^ crossed to Chrimson^A^, polyclonal chicken anti-GFP was used as primary antibody (Thermo Fisher Scientific, AB_2534023), diluted 1:1000 in 0.5 % PBST for neuronal labelling, and a monoclonal mouse anti-Bruchpilot (nc82) antibody (Developmental Studies Hybridoma Bank, AB_2314866), diluted 1:500 in 0.5 % PBST to label neuropil. A polyclonal goat anti-chicken Alexa488 (Thermo Fisher Scientific, AB_2576217) and a polyclonal goat anti-mouse Alex568 (Thermo Fisher Scientific, AB_2534072) were used as secondary antibodies, both diluted 1:500 in 0.5 % PBST.

For Moonwalker^B^ and MDN1^B^ crossed to Chrimson^D^, a polyclonal rabbit anti-dsRed was used as primary antibody (CloneTech, AB_10013483), diluted 1:500 in 0.5 % PBST for neuronal labelling, and a monoclonal mouse anti-Bruchpilot (nc82) antibody (Developmental Studies Hybridoma Bank, AB_2314866), diluted 1:500 in 0.5 % PBST for neuropil labelling. A polyclonal goat anti-rabbit Alex568 (Thermo Fisher Scientific, AB_10563566) and a polyclonal goat anti-mouse Alex647 (Thermo Fisher Scientific, AB_141725) were used as secondary antibodies, diluted 1:500 in 0.5 % PBST.

### EM reconstruction

Renderings of the MDNs in the brain and all their pre- and postsynaptic sites (Fig. 1j, 2a; Extended Data Fig. 1b; Supplemental Data Synapse Topology) were displayed using Codex.ai from FAFB/FlyWire-v783 (Zheng et al. 2018, Buhmann et al. 2021, Eckstein et al. 2023, Dorkenwald et al. 2024, Schlegel et al. 2024). LAL-MDN synapses (Fig. 2e; Supplemental Data Synapse Topology) and MBON-LAL synapses (Supplemental Data Synapse Topology) were displayed using a custom Python script to interact with the FlyWire dataset through the CAVEclient Python package (Dorkenwald et al. 2024).

Renderings of MDN descending arbours in the ventral nerve cord and all their pre- and post-synaptic sites (Fig. 1j, 2a; Extended Data Fig. 1b; Supplemental Data Synapse Topology) were displayed using a custom Python script to interact with the FANC dataset through the FANC Python package (Phelps et al. 2021, Azevedo et al. 2024).

### Video recording of fly movement in circular arena

Experimental procedures followed Bidaye et al. (2020). 1-3-day-old flies of the indicated genotypes and drug treatment were individually placed on a circular arena and given five 10-s light stimulations for optogenetic activation, each pulsed at 50 Hz and 5 ms pulse width, at 50 s intervals. A single trial was considered to consist of a 10 s period with stimulation-light off followed by the 10 s of light stimulation. For the experiments shown in Fig. 1i and Extended Data Fig. 2a, light of 530 nm and 0.65 mW/mm^2^ was used for optogenetic stimulation, whereas for the experiments shown in Extended Data 2b and Extended Data 2c light of 630 nm and 0.80 mW/mm^2^ was used. Continuous low-intensity light at 530 nm and 0.1 mW/mm^2^, in itself insufficient for optogenetic activation, was used throughout to avoid jumping responses upon stimulation-light onset; in addition, low-intensity infrared light of 850 nm and 0.05 mW/mm^2^ was used for video recording at 1280 x 1024 pixel resolution and 30 fps. Stimulation-light was controlled and synchronized to the camera (FLIR BlackFly-S Camera, FL3-U3-13Y3M-C, Richmond, BC, Canada) using a customized Arduino board. Movement of individual flies was tracked using FlyTracker software (Eyjolfsdottir et al. 2014), and data were analysed and plotted in Matlab for translational velocity (mm/s).

### Connectome analyses

Analyses of connectivity from MBONs to MDNs (Fig. 2b,c,d) were performed using the Fly-Wire-v783 dataset with a custom Python script (Dorkenwald et al. 2024, Schlegel et al. 2024). Let *A*_ij_ be the number of synapses onto postsynaptic neuron *i* from presynaptic neuron *j*, thresholded at 10 synapses per entry. We define the normalized weight matrix *W_ij_=A_ij_/Σ_k_A_ik_*, so that each row of *W* sums to 1, and the *n*-step influence from neuron *j* to neuron *i* as *(W^n^)_ij_*. Fig. 2c shows *W*_ij_ as a percentage for the 6 MBON types with the most influence on the MDNs, averaged across MDNs and within MBON type. Fig. 2d shows the percent input from an MBON to MDN via the indicated LAL type, which is calculated as the ratio between the 2-step influence from MBON to MDN without that LAL and the 2-step influence with all intermediaries. The same MBONs, LALs and averaging as in Fig. 2c are used. Connections via the intermediaries that have a large MBON-to-MDN influence are shown.

No MDN-to-DAN pathways with 10 or fewer steps exist where MDNs comprise more than 0.01 percent of the input to a given DAN.

### Modelling

In the model (Fig. 5a-d,f,g), KCs synapse onto MBONs with activities *m*^ap^ and *m*^av^ that promote approach and avoidance, respectively. In the following, we suppress time indices unless they are required for clarity. MBON activities are determined by 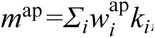, and similarly for *m*^av^, where *k_i_* is the activity of KC*i*, and *w_i_*^ap^ and *w_i_*^av^ are its weights onto MBONs. Odours activate random non-overlapping subsets of *N* KCs. The firing rate of active KCs is *1/N*, and the firing rate of inactive KCs is 0.

The activities of punishment- and reward-responsive DANs *d*^p^ and *d*^r^ at timestep *t* are given by:

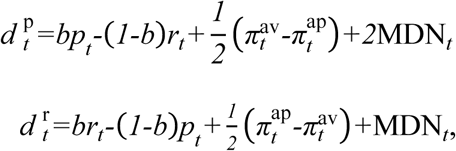

where *b=0.8*, and *p≥0* and *r≥0* represent external punishment and reward. The MDN terms model MDN feedback and influence both DANs, but with a larger magnitude for the punishment-responsive DAN, as empirically observed (Fig. 3e). The prediction components *π*^av^ and *π*^ap^ are calculated as 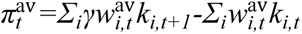, and similarly for *π*^ap^, where *γ=0.99* (Fig. 5c,d) or *γ=0.8* (Fig. 5f,g) is the temporal discount factor.

KC-to-MBON weights are updated as 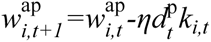 and 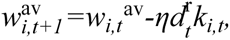 where *η=0.1* is the learning rate. After this update, at each timestep the weights are subtractively normalized such that 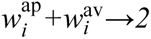 asymptotically: 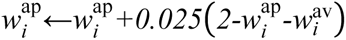 and 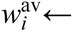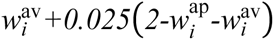 (Extended Data Fig. 9h,i).

The difference *V=m*^ap^*-m*^av^ represents the model’s estimate of the current odour’s value and controls its approach and avoidance behaviour. At each timepoint, the model fly approaches the odour source, stays in place, or avoids the odour source with probabilities [*p*_approach_, *p*_stay_, *p*_avoid_]. When *V>0.01*, these probabilities are [*0.8, 0.1, 0.1*]. When *V<-0.01*, they are [*0.1, 0.1, 0.8*]. Otherwise, they are [*0.4, 0.2, 0.4*].

For models with MDN feedback, MDN*_t_=0.1* if, on the previous timestep, the “avoid” action was selected and *V<-0.001*. Otherwise, MDN*_t_=0.* This value-gating condition improves model performance by not reinforcing random “avoid” actions (Extended Data Fig. 9d-g), although our qualitative results do not depend on this choice. For models without MDN feedback, MDN*=0* and the model is equivalent to temporal difference learning (Sutton & Barto 2018).

Extinction experiments (Fig. 5e-g) consist of two phases – one training phase and one extinction phase – each with 35 timesteps, followed by a test phase. During the training phase, an odour is presented and an external punishment of magnitude -0.4 is delivered every three timesteps, starting on the second timestep. During the extinction phase, odour is either present or absent on each side of the arena according to the indicated protocol (Fig. 5e), with no external reinforcement. During the test phase, each model’s probability of choosing the punishment-predicting odour is given by a softmax function 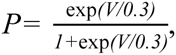, where *V* is the final estimated value of the punished odour following timestep 72. After each phase, a timestep in which no odour is present occurs.

### Statistical analyses

Non-parametric statistical tests were used throughout (Statistica 11.0, StatSoft Hamburg, Germany; Python package Pingouin 0.5.4; R 2.15.1, http://www.r-project.org). The Kruskal-Wallis test (KW test) was applied for comparisons between more than two groups. For subsequent two-group comparisons, or for cases with only pairwise between-group comparisons, the Mann-Whitney U-test (MW-U test) was performed. To test whether the values of a given group differed from chance levels, i.e. from zero, the one-sample sign test (OSS test) was used. Within-animal comparisons used the Wilcoxon signed-rank test. Significance levels of multiple tests were adjusted by a Bonferroni-Holm correction to keep the experiment-wide type 1 error limited to 0.05 (Holm 1979). Data are presented as box plots with the median shown as the middle line and the interquartile range (box) and 10th/90th percentiles as box boundaries and whiskers, respectively. In cases of within-animal comparisons, data of individual flies are displayed and connected across repeated measurements. For conditioning and choice experiments, a sample size of N= 1 included ∼60-100 flies; video recordings of fly movement were done on the indicated number of individual flies. For model fly extinction experiments, a sample size of N= 1 included 20 individual model flies. For two-photon calcium imaging data, the ΔΔF/F_0_ values were used for all statistical analyses on the indicated number N of individual flies. The data in Extended Data Fig. 5xref>b’’,c’’ were analysed using the Friedman test.

## Acknowledgements

The authors thank Benjamin Bargeron and Gianna Vitelli for help with EM reconstruction; Hongbo Jia and the LIN-CNI facility for help with setting up fly *in vivo* imaging; Davide Raccuglia for help with brain explant imaging; Ansgar Büschges for drawing our attention to the role of action-valence feedback during extinction protocols; Marta Andreatta, Paul Pauli, Marcella Woud, Till Bockemühl and Ansgar Büschges for comments on and Rupert D.V. Glasgow for corrections of earlier drafts of the manuscript. The authors acknowledge support from the Kavli Foundation (JTS, AL-K); the Gatsby Charitable Foundation *GAT3708* (JTS, AL-K); the National Science Foundation: Graduate Research Fellowship *DGE-2036197* (JTS); the Deutsche Forschungsgemeinschaft (DFG): *FOR 2705* (DO, IGK, BG), *TRR 265* (DO), Walter Benjamin Fellowship *MA10161/1-1* (NM); the Max-Planck-Society (S.S.B.); the European Research Council: ERC Consolidator Grant *101088502* (DO); the Deutscher Akademischer Austauschdienst: Research Grant Doctoral Program *57552340* (UM); the Ministry of Culture and Science North Rhine-Westphalia: iBehave *NW21-049* (IGK); the European Union: Erasmus+ *2020-1-PL01-KA103-078995* (AP); The Company of Biologists: Travelling Fellowship *JEBTF2003429* (AP); the National Institutes of Health: Award *RF1DA06722* (AL-K); the DFG: *Cluster of Excellence Multiscale Bioimaging* and the Ministry of Science and Culture Lower Saxony: *zukunft.niedersachsen* (OB), as well as the Leibniz Institute for Neurobiology (BG).

## Author contributions

F.A., J.T.S., A.L.-K., C.K. and B.G. conceived of the study and designed the experiments. F.A., C.K., A.P. and U.M. performed conditioning and choice experiments supervised by B.G. and I.C.G.K.. F.A. performed *in vivo* calcium imaging experiments supervised by O.B.. J.T.S. and A.L.-K. performed connectome analyses and computational modelling. N.M., S.S.B. and K.M. performed immunohistochemistry, EM rendering, and fly locomotion tracking. M.-M.H. performed explant calcium imaging supervised by D.O.. F.A., J.T.S., B.G., O.B. and C.K. analysed and interpreted the data, and visualized the results with input from A.L.-K., S.S.B., N.M., D.O., M.-M.H; U.M., I.C.G.K., K.M., and A.P.. F.A., J.T.S. and B.G. wrote the original manuscript with reviewing and editing input from A.L.-K., O.B., C.K., D.O. and I.C.G.K.. B.G. supervised the study.

## Declaration of interests

The authors declare no competing interests.

---

**Extended Data Fig. 1.**
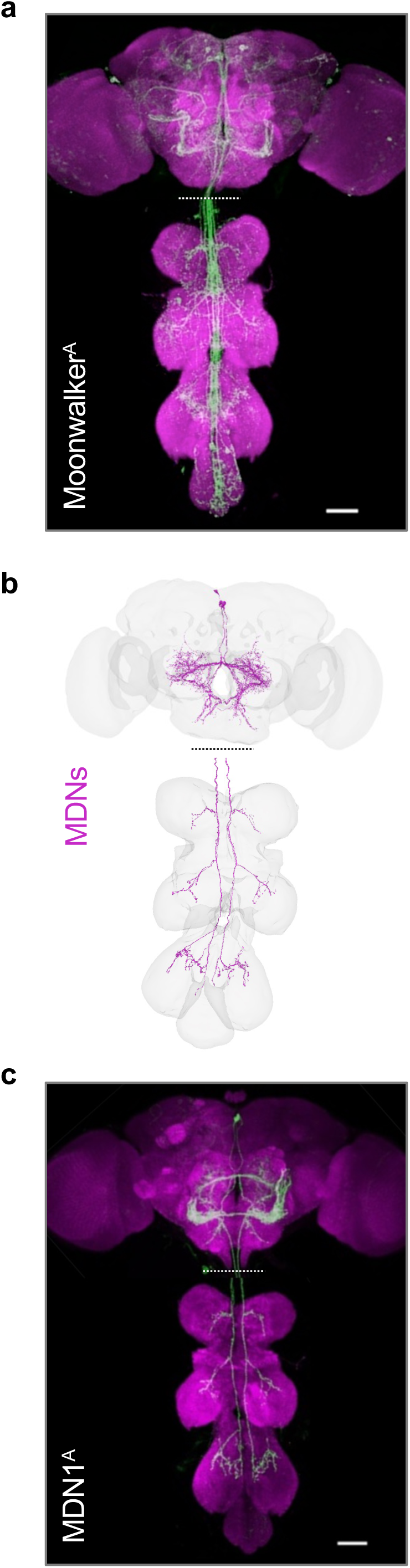
Expression patterns of Gal4 drivers and MDN reconstruction. **a-c,** Higher-resolution images of the anatomy panels in Fig. 1a **(a)** and Fig. 1j **(b,c)**. Anti-GFP labelling (green) driven by Moonwalker^A^ **(a)** and MDN1^A^ **(c)** is shown along with neuropil labelled by anti-Bruchpilot (magenta). Shown in **(b)** is the EM reconstruction of the MDN neurons (magenta) in the context of the neuropil (grey mesh). Genotypes: Moonwalker^A^>Chrimson^A^ **(a)**, MDN^A^>Chrimson^A^ **(c)**.

**Extended Data Fig. 2.**
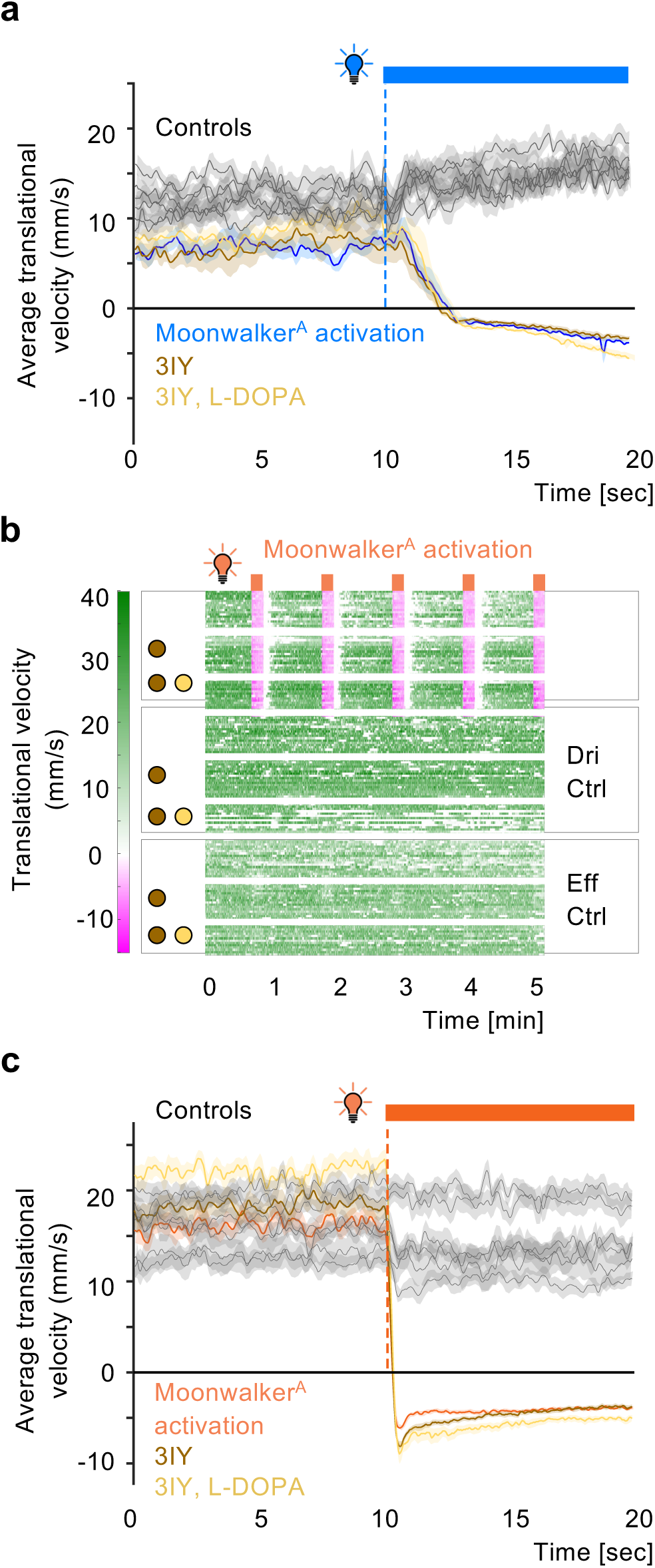
Optogenetically induced backward movement is unaffected by 3IY. **a,** For the experiment shown in Fig. 1i, translational velocity (mm/s) was averaged across trials for each fly, for the 10 s before and during 10-s optogenetic activation (blue bar), and is plotted as mean ± SEM. For the experimental genotype (Moonwalker^A^>ChR2XXL^A^, coloured traces), optogenetic activation leads to negative translational velocity, i.e. backward movement, regardless of the indicated drug treatment (brown: 3IY, light brown: additional supply of L-DOPA). In genetic controls (Moonwalker^A^>+, +>ChR2XXL^A^) no backward movement is observed, likewise regardless of drug treatment (black traces) (N= 12,8,12,16,12,12,12,12,12). **b,** As in Fig. 1i, for Chrimson as the effector, showing that 3IY does not impact moonwalker-induced movement for an independent effector. Experimental genotype; Moonwalker^A^>Chrimson^A^; genetic controls: Moonwalker^A^>+ (Dri Ctrl), +>Chrimson^A^ (Eff Ctrl) (N= 16,16,12,16,16,12,16,16,12). **c,** as in **(a)**, for the experiment shown in **(b)**.

**Extended Data Fig. 3.**
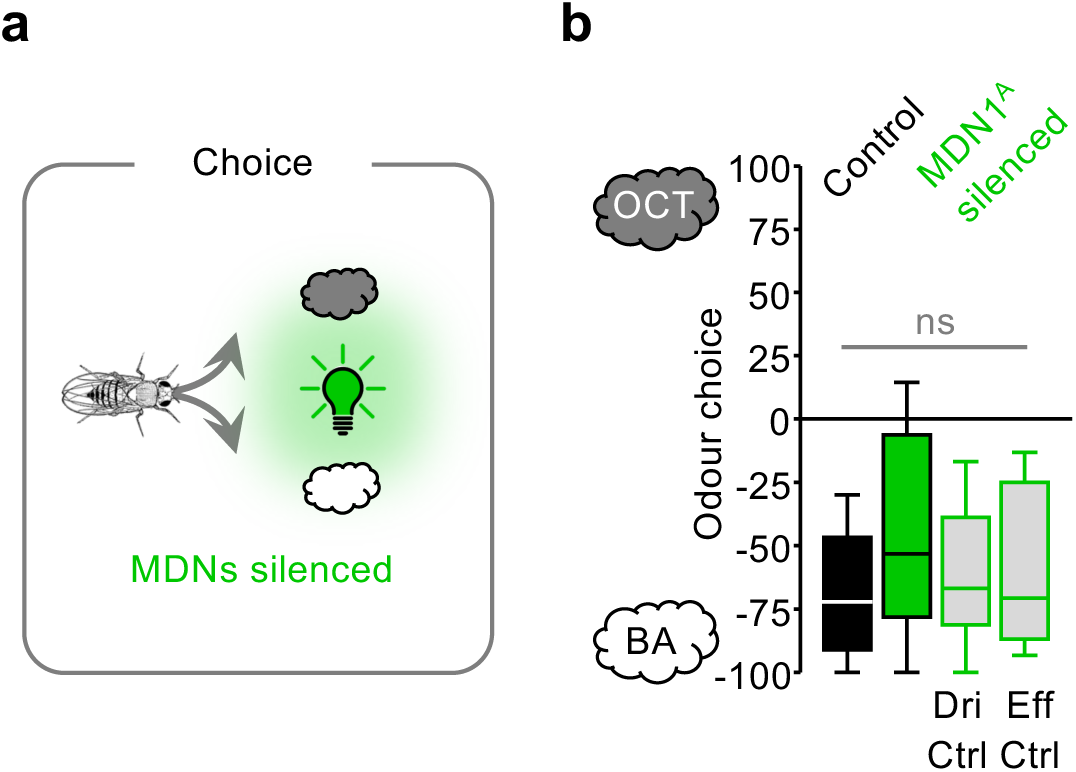
Innate olfactory choice behaviour is unaffected by MDN silencing. **a,b,** Overview **(a)** and outcome **(b)** of innate olfactory choice experiments. Clouds: odours. Light bulb: optogenetic silencing of MDNs. Odour choice in experimentally naïve flies does not differ for the experimental genotype (MDN1^A^>GtACR1) under control conditions without light stimulation (black) from the MDN1^A^-silenced condition (green), or from genetic controls under light stimulation (grey) (Dri Ctrl: MDN1^A^>+, Eff Ctrl: +>GtACR1) (N= 18,19,23,22). Box-whisker plots show median, interquartile range (box) and 10th/90th percentiles (whiskers). Data were analysed across groups by a Kruskal-Wallis test (ns: P> 0.05).

**Extended Data Fig. 4.**
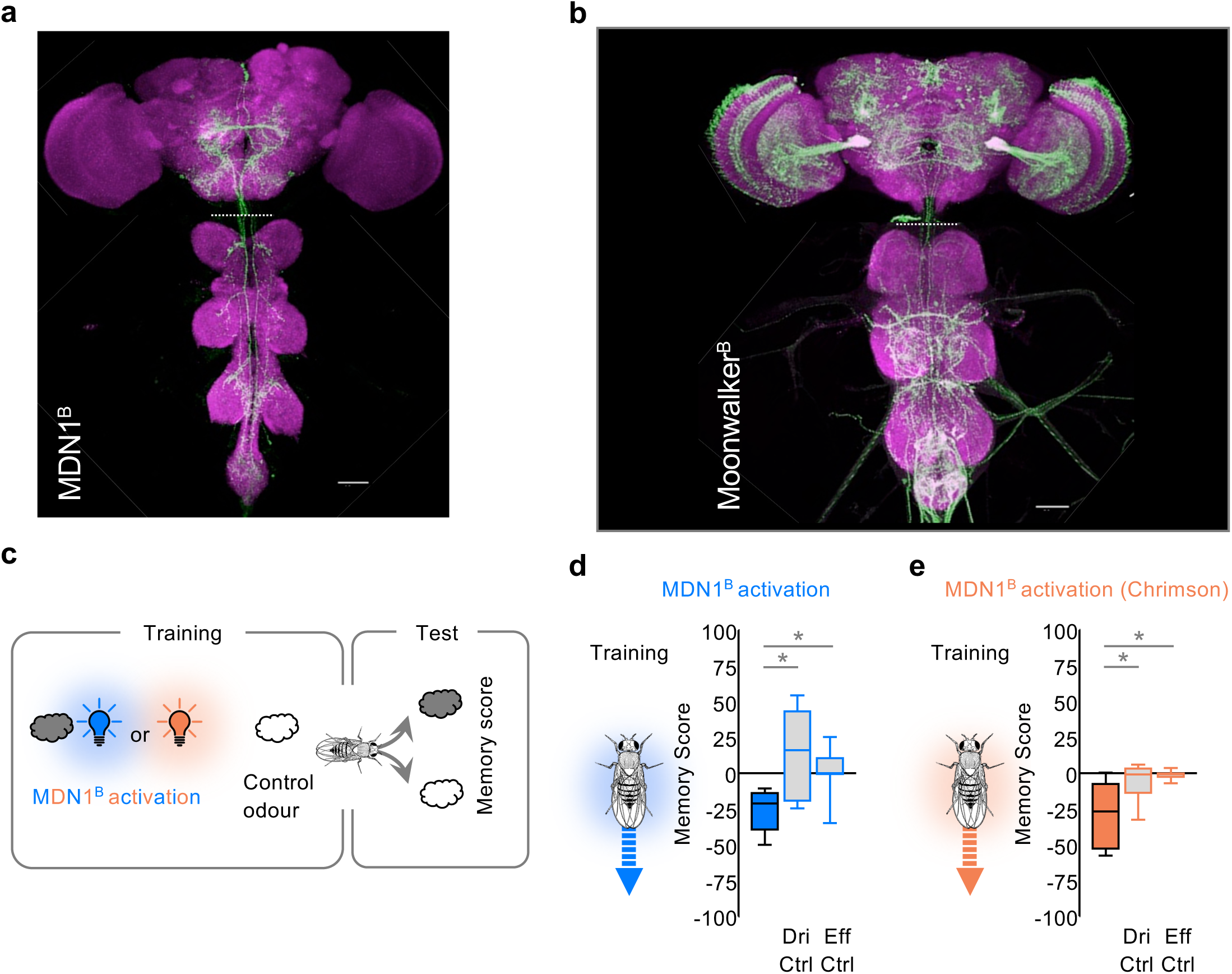
Characterization of lexA drivers. **a,b,** Anti-GFP labelling (green) driven by MDN1^B^ **(a)** and Moonwalker^B^ **(b)** along with neuropil labelled with anti-Bruchpilot (magenta). Other details as in the legend of Fig. 1a,j. **c-e,** Overview **(c)** and outcome of pairing odour with MDN1^B^ activation using either ChR2XXL (blue: MDN1^B^>ChR2XXL^B^) **(d)**, or Chrimson (orange: MDN1^B^>Chrimson^B^) **(e)**, resulting in aversive memory in the experimental genotypes but not in genetic controls (Dri Ctrl: MDN1^B^>+, Eff Ctrl: +>ChR2XXL^B^ **(d)** or +>Chrimson^B^ **(e)**) (N= 12,12,10; 12,10,11). Box-whisker plots show median, interquartile range (box) and 10th/90th percentiles (whiskers). Data were analysed across groups by Kruskal-Wallis tests (P< 0.05), followed by pairwise comparisons (Mann-Whitney U-tests, *P< 0.05 with Bonferroni-Holm correction). Other details as in the legend of Fig. 1j.

**Extended Data Fig. 5.**
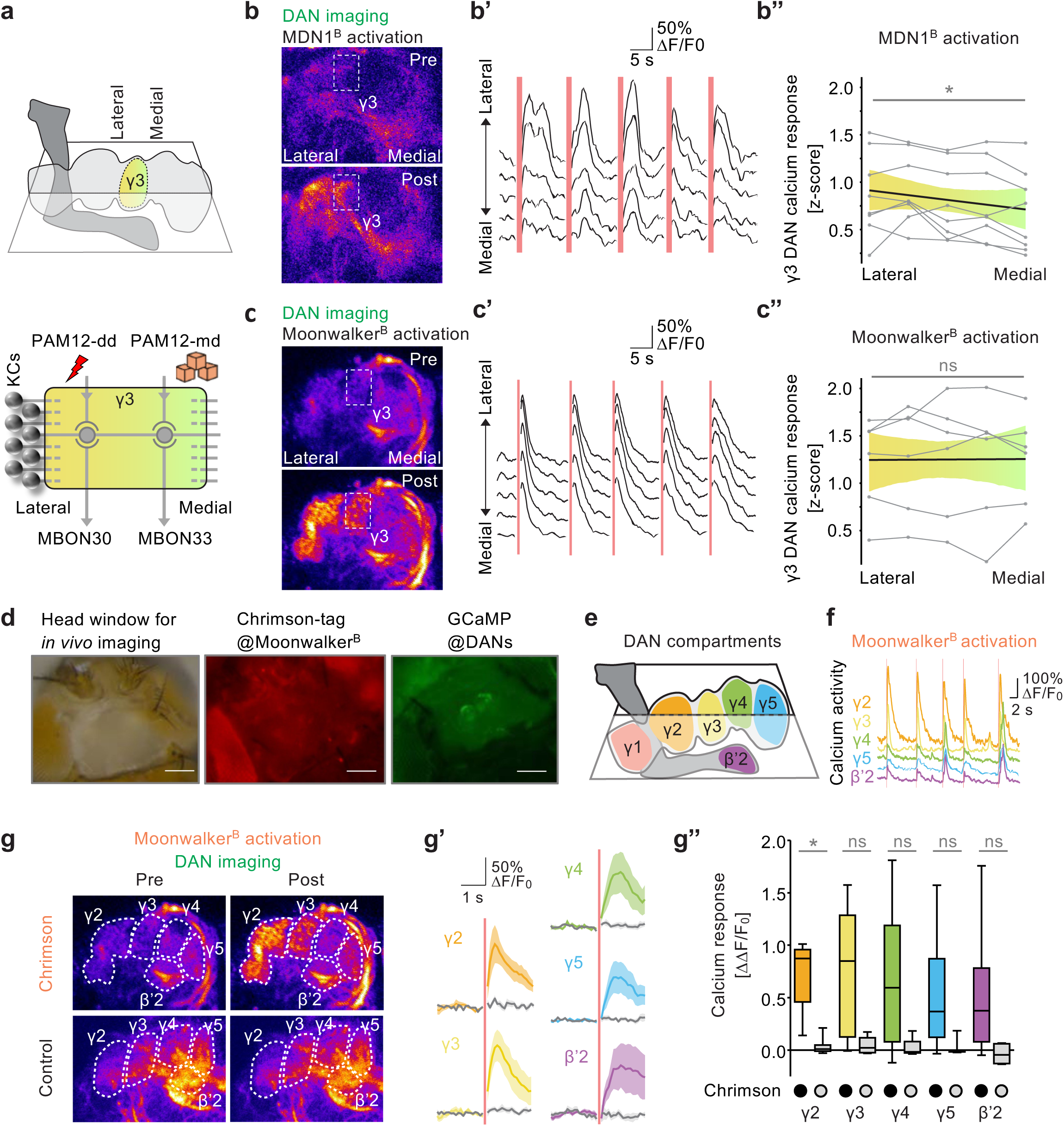
Compartmental topology of DAN engagement by MDN and moonwalker neuron activation. **a,** Imaging plane and topology of the γ3 compartment (top) and its internal organization (Li et al. 2020) (bottom). Other KCs connect only to PAM12-dd or PAM12-md (not shown). **b-b’’,** *In vivo* calcium imaging of flies expressing GCaMP across the DANs and optogenetic activation of MDNs by 200-ms red light stimulation, re-analysed from Fig. 3e to reveal within-γ3 topology (genotype: MDN1^B^>Chrimson^C^; DANs>GCaMP). Shown in **(b)** are average intensity projections of a sample recording 2 s before/ after stimulation (Pre/ Post). For the γ3 compartment region of interest (stippled rectangle in **b**), calcium transients (ΔF/F_0_) for spatial bins from lateral (top) to medial (bottom) are displayed for consecutive activation trials (red vertical bars) **(b’)**. Z-scored average calcium responses show stronger DAN engagement in lateral than medial bins (N = 8) **(b’’)**. **c-c’’,** As in **(b-b’’)**, for 20-ms activation of the full set of moonwalker neurons (experimental genotype: Moonwalker^B^>Chrimson^C^; DANs>GCaMP), suggesting uniform DAN activation throughout γ3 (N= 6). **d-g’’,** Analysis of the experiment in **(c)** for all imaged compartments. For a fly of the experimental genotype with the head capsule opened for imaging, panel **(d)** shows brightfield (left) and fluorescence microscopy images to confirm transgene expression (middle and right). Scale bar 100 µm. For the compartments covered at the chosen imaging plane **(e)**, sample traces of raw calcium transients are shown for 5 trials of activation (red vertical lines) **(f)**. Panel **(g)** as in **(c)**, for the experimental genotype (top) and Controls (Moonwalker^B^>+; DANs>GCaMP) (bottom). Calcium transients from the indicated compartments upon activation in the experimental genotype (coloured traces) and in Controls (grey traces) **(g’)**. Activation of moonwalker neurons results in significant calcium responses (ΔΔF/F_0_) of DANs in only the γ2 compartment (N= 6,6) **(g’’)**. Box-whisker plots show median, interquartile range (box) and 10th/90th percentiles (whiskers). Calcium transients are plotted as mean ± SEM. Data were analysed by Friedman tests (*P< 0.05, ns: P> 0.05) **(b’’, c’’)** or by a Kruskal-Wallis test (P< 0.05), followed by pairwise comparisons (Mann-Whitney U-tests, *P< 0.05 with Bonferroni-Holm correction) **(g’’)**. **(b’’, c’’, g’, g’’)** are based on the first optogenetic activation trial. Additional information in Extended Data Fig. 4b.

**Extended Data Fig. 6.**
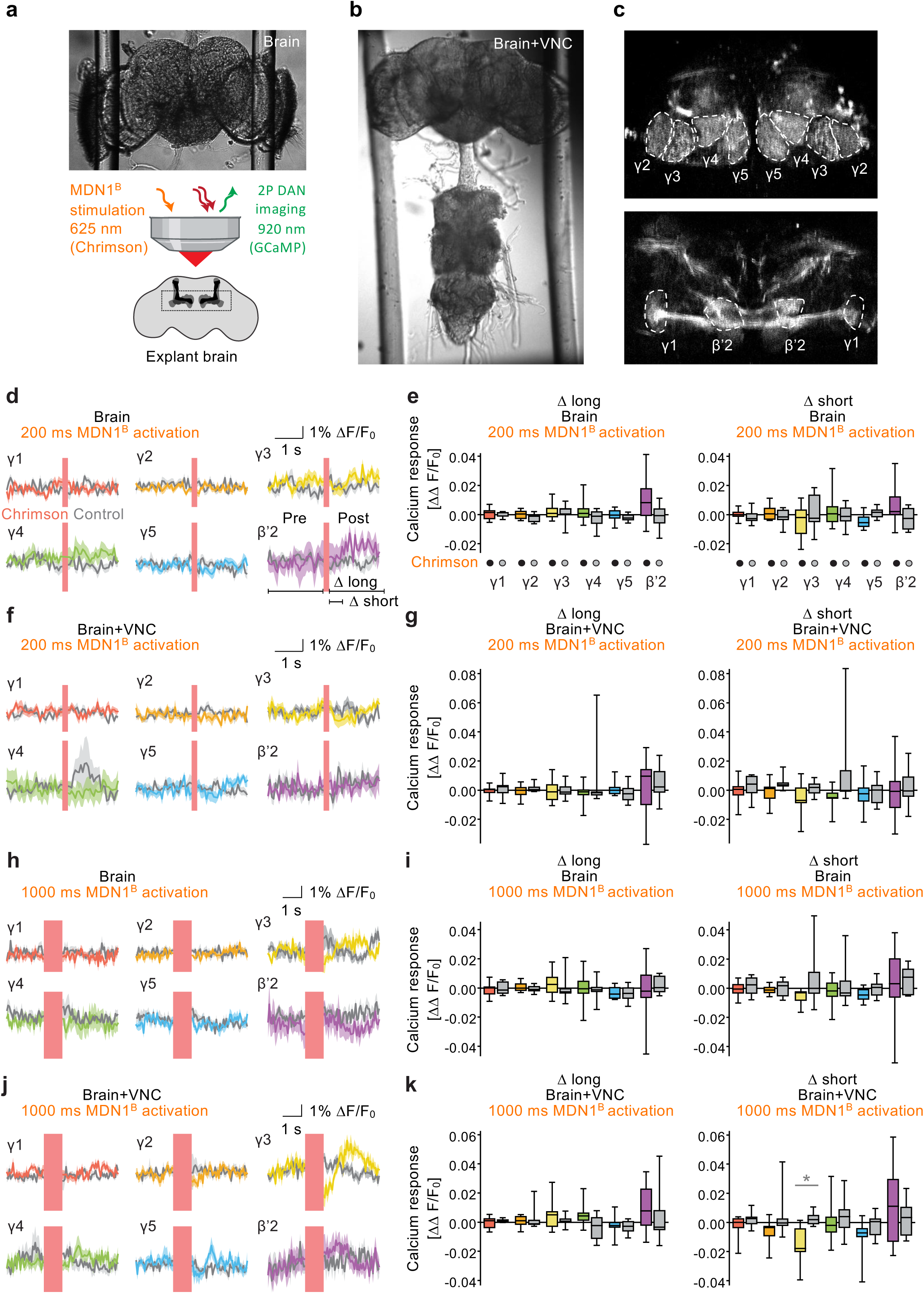
Activation of MDNs in explant preparations does not engage DANs. **a,b,** Overview of imaging setup of explant brain (**a**) and brain-plus-VNC (**b**) preparation. MDN1^B^ was activated using Chrimson with 200 ms or 1000 ms red light while calcium transients were monitored across the DANs of the horizontal lobe compartments by GCaMP6f. **c,** Average intensity projections from representative brain preparations of the experimental genotype (MDN1^B^>Chrimson^C^; DANs>GCaMP) under two-photon illumination, with a focus on γ2− γ5 DANs (top) and γ1 and β’2 DANs (bottom). Dashed lines indicate compartment boundaries. **d-e,** Calcium transients (ΔF/F_0_) from 2 s before to 2 s after MDN1^B^ activation with 200 ms of red light (red vertical bar) in brains of the experimental genotype (coloured traces; N= 10) or control brains (grey traces, N= 10) (MDN1^B^>+; DANs>GCaMP) **(d)**. Calcium responses (ΔΔF/F_0_) in these brains comparing 2 s before activation to 2 s after activation (Δ long) or to 0.5 s after activation (Δ short). No significant differences between Chrimson-expressing and control brains were observed. **f,g,** As in **(d-e)**, for brain-plus-VNC preparations (N= 10,10). **h-k**, As in **(d-g)**, for 1000-ms red light, revealing a decrease in calcium responses in γ3 DANs of brain-plus-VNC preparations after 1000 ms stimulation and for the Δ short period, arguing for the functionality of the assay. Box-whisker plots show median, interquartile range (box) and 10th/90th percentiles (whiskers). Calcium transients are plotted as mean ± SEM. Data were analysed by Kruskal-Wallis tests (P< 0.05), followed by pairwise comparisons (Mann-Whitney U-tests, *P< 0.05 with Bonferroni-Holm correction).

**Extended Data Fig. 7.**
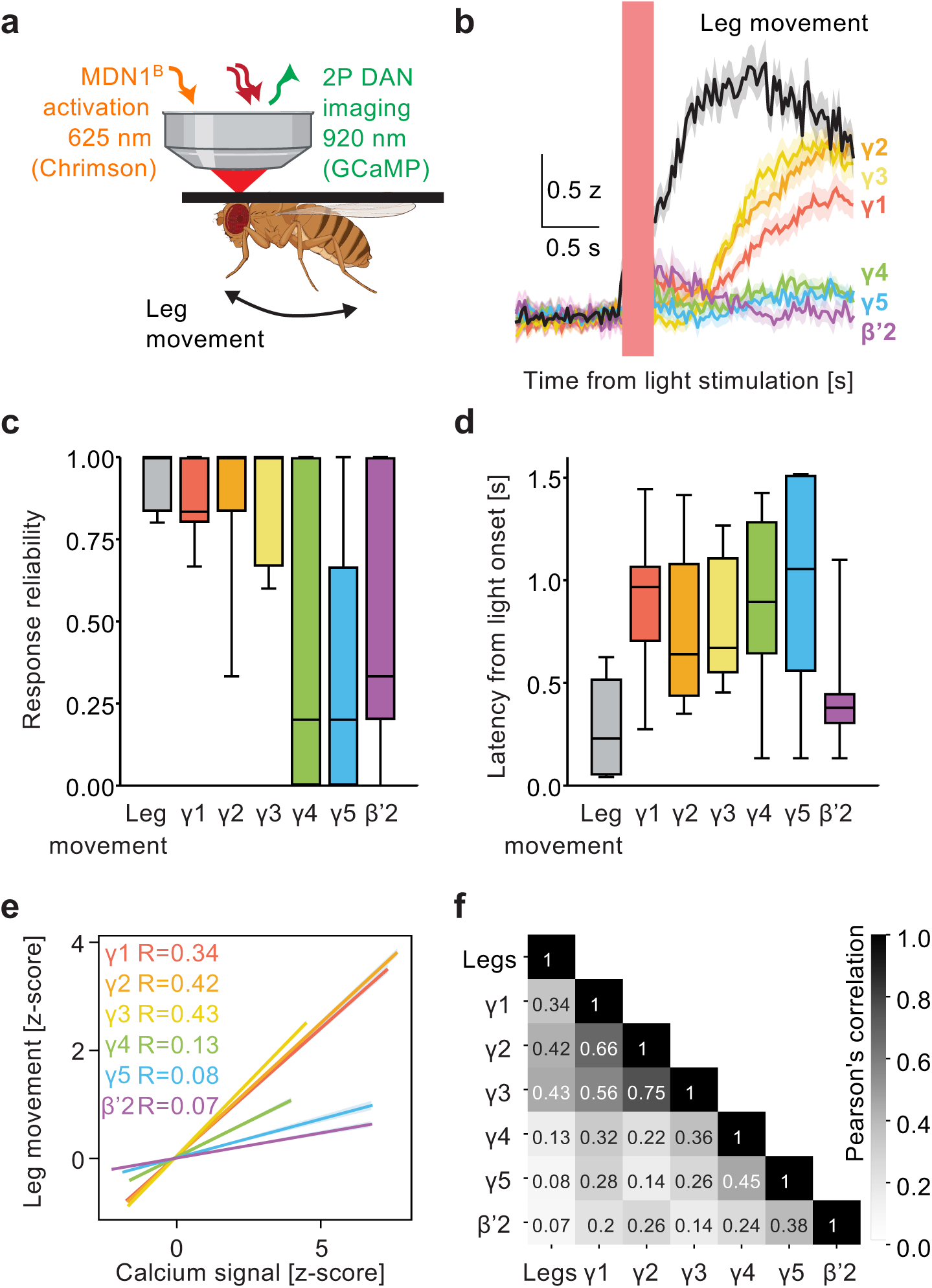
Temporal relationship of MDN-evoked leg movement and DAN engagement. **a,** Sketch of the *in vivo* imaging setup. MDN1^B^ was activated using Chrimson with 200-ms red light stimuli while calcium transients were monitored in horizontal lobe DANs. Animals were free to move their legs. Genotype: MDN1^B^>Chrimson^C^; DANs>GCaMP. **b,** Average calcium signals (coloured traces) and leg movement (black traces) around the time of light stimulation (red vertical bar). Shown are z-scored values; leg movement refers to the motion energy calculated from the legs. **c,d,** Response reliability **(c)** and latency **(d)** of leg movements (grey) and DAN engagement (colour-coded by compartment) upon light stimulation. Responses are defined as values >2 standard deviations above the pre-2 s average within a 2 s time window after light stimulation. Reliability refers to the probability of responses across the five light stimulations; latency refers to the first time the response threshold is crossed after light stimulation. **e,f,** Pearson’s correlation of leg movement and engagement of the indicated DANs **(e)** and corresponding pairwise correlations **(f)**. Panels show analyses of data from Fig. 3e and Fig. 4a-e (before trapping) for which reliable quantifications of leg movement were possible across all light stimulation trials (N= 11) **(b,c, e,f)** based on individual time data points **(e,f)**; for the response latency measure in **(d)** sample sizes are naturally lower for the compartments in which no responses were observed (N= 11,11,10,11,6,6,9). Box-whisker plots show median, interquartile range (box) and 10th/90th percentiles (whiskers). Calcium signals are plotted as mean ± SEM.

**Extended Data Fig. 8.**
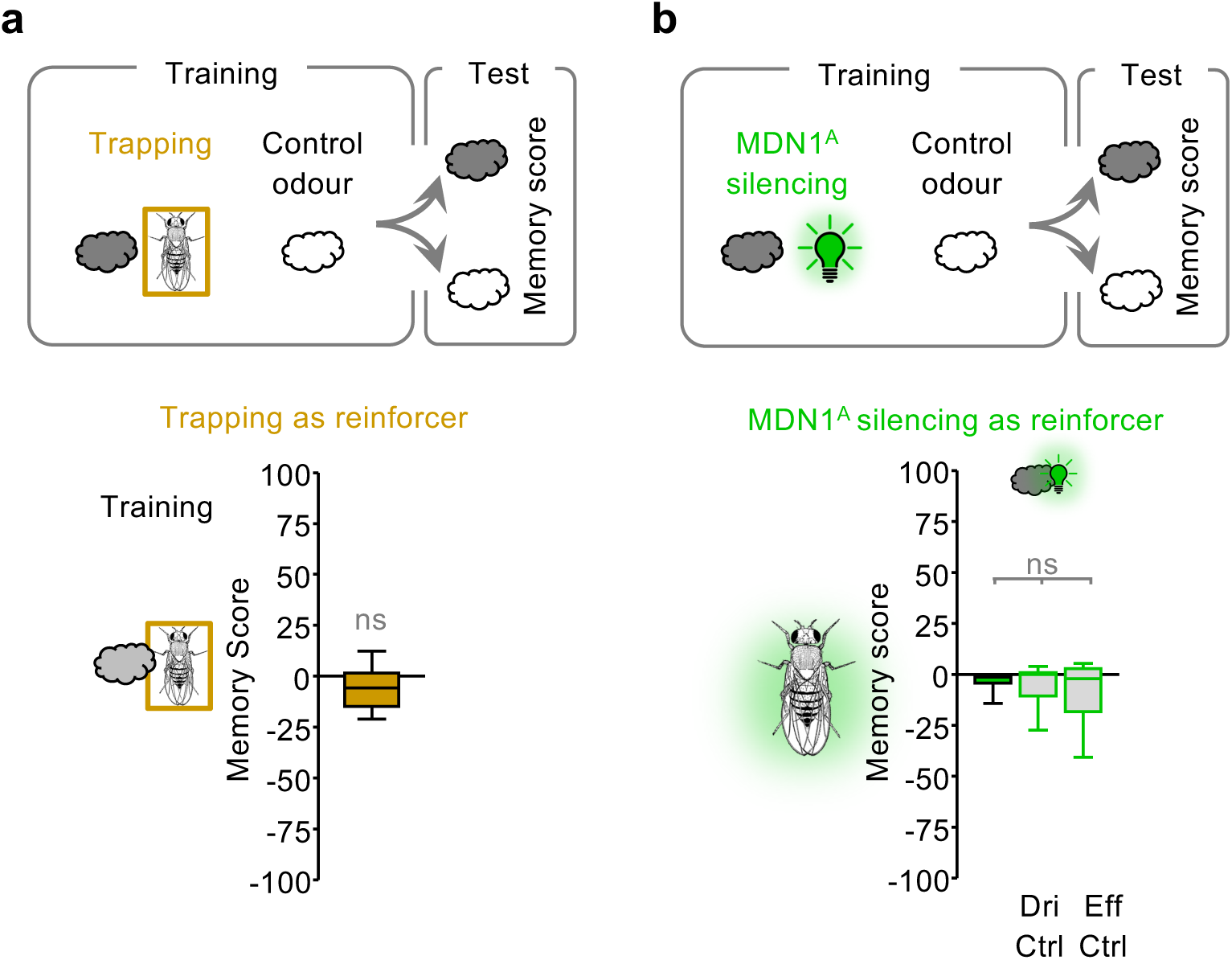
No punishing effect of restraint or of MDN silencing. **a,** Overview and outcome of olfactory learning experiments with movement restraint (Trapping; also see Fig. 4f, top) as reinforcer (genotype: CantonS, N= 13), suggesting that it has no punishing effect. Clouds: odours. Orange box: movement restraint. **b,** Overview and outcome of olfactory learning experiments with MDN silencing as reinforcer, suggesting that it has no punishing effect. Clouds: odours. Light bulb: optogenetic silencing of MDNs (MDN1^A^). Experimental genotype: MDN1^A^>GtACR1, Dri Ctrl: MDN1^A^>+, Eff Ctrl: +>GtACR1 (N= 12,12,12). Box-whisker plots show median, interquartile range (box) and 10th/90th percentiles (whiskers). Data were analysed with one-sample sign (OSS) **(a)** and Kruskal-Wallis tests **(b)** (ns: P> 0.05). Additional information in Supplemental Video 4.

**Extended Data Fig. 9.**
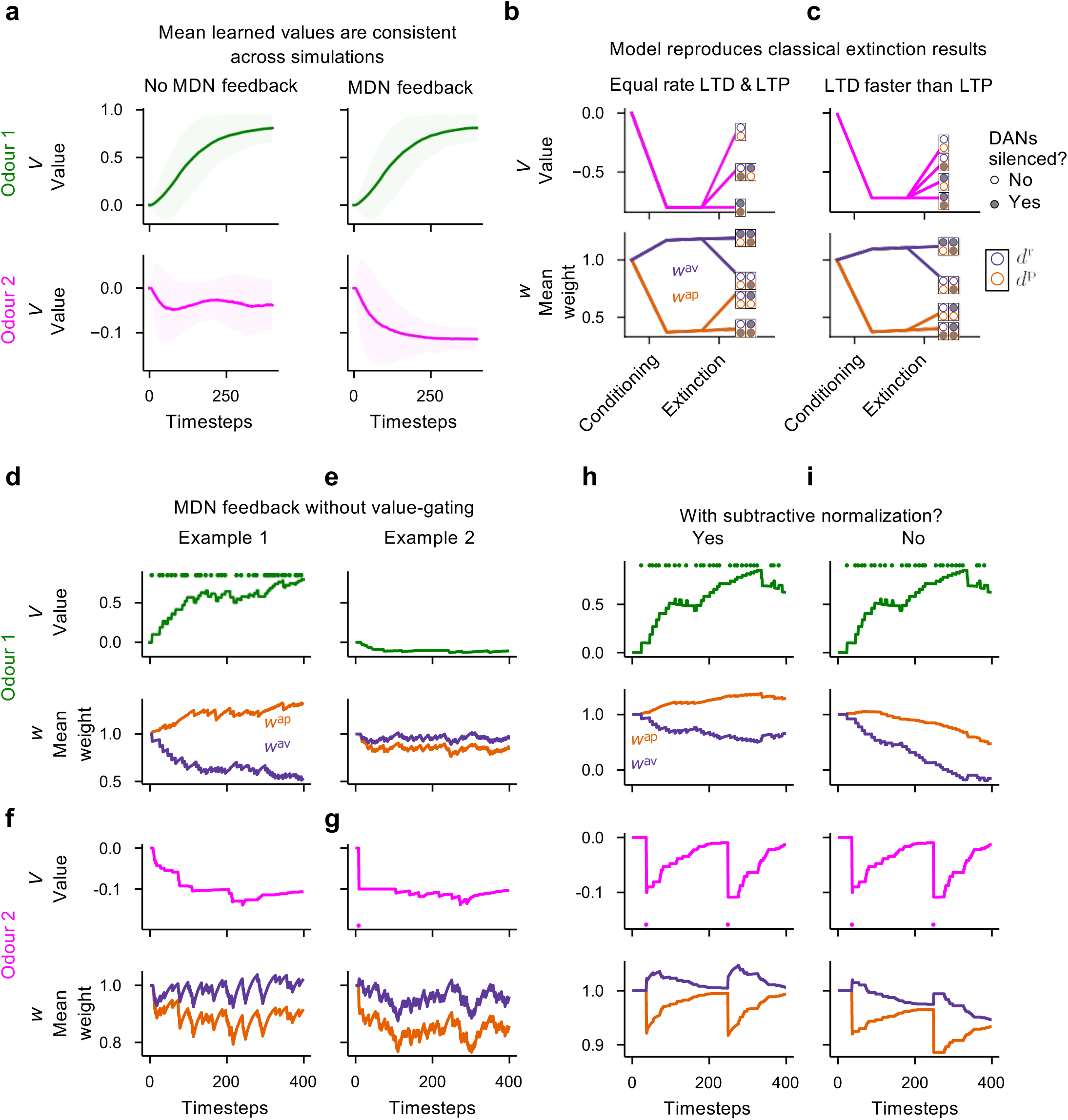
Effects of varying model assumptions. **a,** Mean learned odour values across 400 model flies, showing that the effect in Fig. 5d is consistent. Shaded regions show ± 2 standard deviations. **b,** A model fly analogous to the ones shown in Fig. 5a,b,f responding to a classical extinction task, with DANs intact or silenced during extinction. A model fly is exposed to odour and then shock during conditioning and learns the association value *V* (pink line). It is then exposed to odour without shock and extinguishes the association. Conditioning mainly induces depression in the approach compartment weights *w*^ap^ (orange line). Extinction effects are shared between compartments. **c,** As in **(b)**, but with LTP at a slower rate, whereas LTD remains the same. This tweak to the model, which does not affect the qualitative results presented in Fig. 5, makes the dynamics of learning and extinction consistent with previous work (Felsenberg et al. 2018). Extinction occurs mainly via the ‘avoid’ compartment weights *w*^av^ (purple line), and silencing reward DANs leads to loss of extinction (*V*), whereas silencing punishment DANs has a smaller effect on extinction (*V*). **d-g,** Model with MDN feedback that is not gated by value for odour 1 **(d-e)** or odour 2 **(f,g)** in two example model flies (**d,f** and **e,g**). In **(e)**, the model fly spontaneously walks backwards in the presence of odour 1 and so learns an aversive association with it, missing out on the reward. In **(f)**, the model fly learns an aversive association with odour 2 by repeatedly avoiding but without ever actually experiencing the negative reinforcement. This figure demonstrates the role of gating feedback by value. **h,** A model fly analogous to the one shown in Fig. 5d for the ‘No MDN feedback’ condition, for comparison with **(i)**, which shows the model without subtractive normalization of the weights. This leads to weights continuing to decrease over time, eventually leading to negative synaptic weights. The learned values (*V*) are identical in **(h,i)** because the difference between the weights (*w*) remains the same.

**Extended Data Fig. 10.**
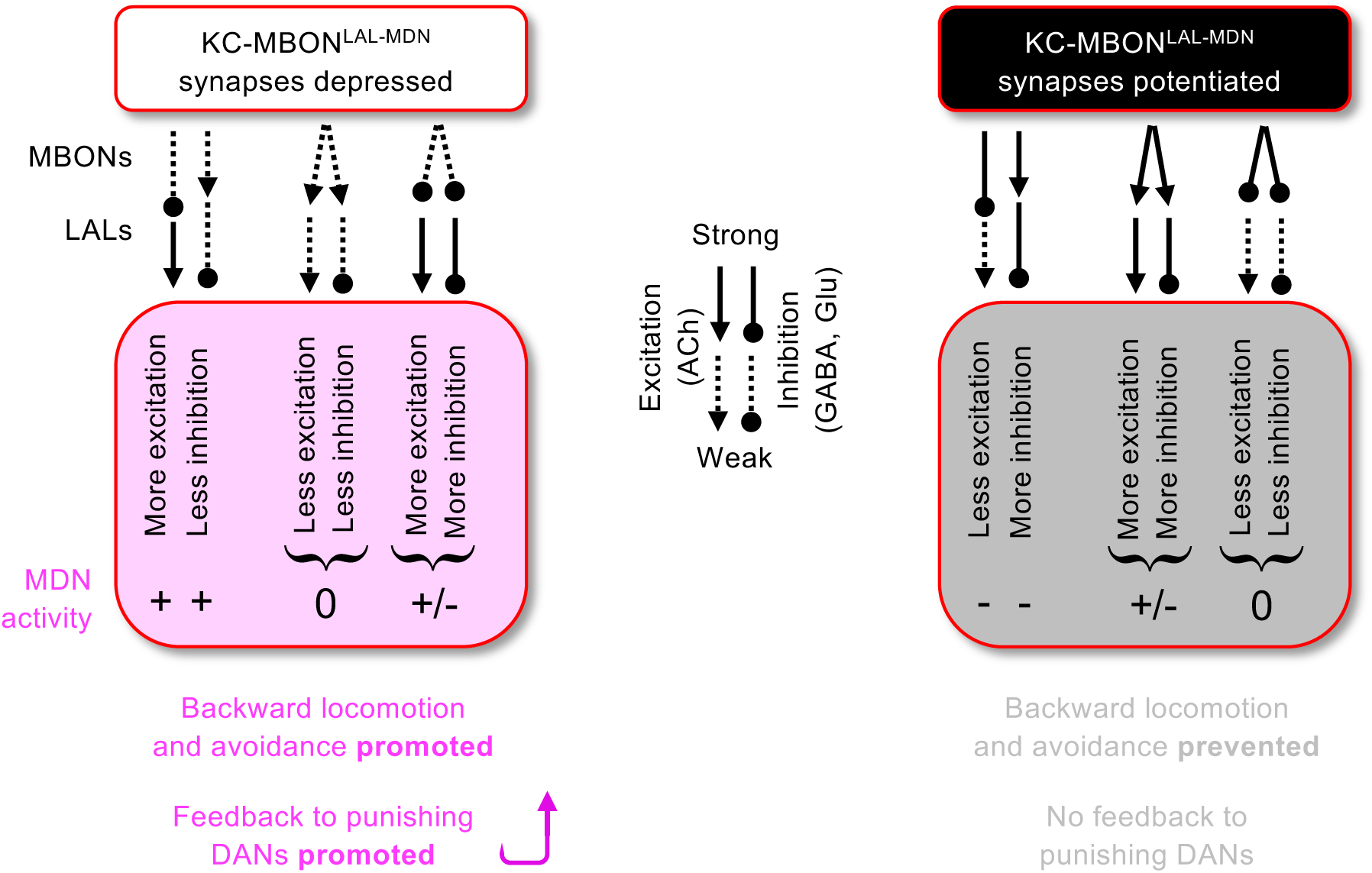
Scenarios of how KC-MBON plasticity affects MDNs. Schematic summary of Fig. 2c. Shown are scenarios of depressed (left) or potentiated (right) synapses between the KCs and the MBONs upstream of MDNs (KC-MBON^LAL-MDN^). Through pathways with sign-inversion (inhibitory MBON to excitatory LAL or vice versa) depressed/ potentiated KC-MBON synapses promote/ prevent MDN activity and avoidance. For MBONs that give rise to parallel pathways with excitatory and inhibitory effects on MDNs the scenario is different. Using synapse number as a proxy for connection strength, these have nearly equal influences on the MDNs (Fig. 2d). Through these MBONs, the MDNs receive both more excitation and more inhibition (originating in inhibitory MBONs when the KC-MBON synapses are depressed, but in excitatory MBONs when they are potentiated, whereas the respectively other MBONs in the indicated extreme case lose their influence on the MDNs). Terminals from LAL171,172 and LAL051 as the main conduits in these pathways are located in close proximity to one another on the MDN dendrites (Fig. 2e; Supplemental Data Synaptic Topology). Nonlinear dendritic interactions based on this proximity may thus render the MDNs particularly sensitive to modulation.

**Supplemental Video 1│ Activating MDNs while imaging DANs.** Flies were stimulated with 5 repetitions of 200 ms red light for MDN activation (red squares) while calcium transients were monitored in horizontal lobe DANs (γ1-γ5 and β’2). First a representative fly of the experimental genotype expressing Chrimson in the MDNs is shown (MDN1^B^>Chrimson^C^; DANs>GCaMP), followed by a control fly not expressing Chrimson (MDN1^B^>+; DANs>GCaMP). Calcium responses are observed in the experimental fly but not in the control fly (quantification in Fig. 3e). The video plays at 3.75X speed.

**Supplemental Video 2│ Activating moonwalker neurons while imaging DANs.** Flies were stimulated with 5 repetitions of 20 ms red light for moonwalker neuron activation (red squares) while calcium transients were monitored in horizontal lobe DANs (γ2-γ5 and β’2). First a representative fly of the experimental genotype expressing Chrimson in the moonwalker neurons is shown (Moonwalker^B^>Chrimson^C^; DANs>GCaMP), followed by a control fly not expressing Chrimson (Moonwalker^B^>+; DANs>GCaMP). Calcium responses are observed in the experimental fly but not in the control fly (quantification in Extended Data Fig. 5c-g’’). The video plays at 6X speed.

**Supplemental Video 3│ Restraining leg movement (trapping) abolishes MDN-evoked DAN activation.** Flies were stimulated with 5 repetitions of 200 ms red light for MDN activation (red squares, red vertical bars) before, while or after being ‘trapped’ by having their leg movements restrained with a piece of cotton wool, while calcium transients were monitored in horizontal lobe DANs (γ1-γ5 and β’2) (MDN1^B^>Chrimson^C^; DANs>GCaMP). Shown are leg movements (top left), calcium transients (top right), and the corresponding traces (bottom) for one and the same representative fly before being trapped, while trapped, and after being trapped. MDN-evoked DAN activity is transiently abolished while the fly is trapped (quantification in Fig. 4e). The video plays at 3.75X speed.

**Supplemental Video 4│ Restraining movement (trapping) during training.** The video shows the procedure for transiently restraining movement (trapping) for cohorts of 40-60 flies. Trapping was performed without apparent harm, as indicated by the natural movement of flies after trapping. Next, a representative example of the trapping procedure is demonstrated in the training apparatus. This shows how odour containers were attached during training as well as how flies were released from trapping before the choice test.

## Notes

### Competing Interest Statement

The authors have declared no competing interest.

