## Supplemental Methods Table 1 for "Avoidance engages dopaminergic punishment in *Drosophila*"

| Simplified genotype | Genotype | Source | Used in Figures |
|---|---|---|---|
| Moonwalker$^{A}$ | VT050660-Gal4 | Bidaye et al. 2014 | Fig. 1a,c-i<br>Ext. Data Fig. 1a, 2 |
| Moonwalker$^{B}$ | VT044845-lexA | Sen et al. 2017 | Ext. Data Fig. 4b, 5c-g |
| MDN1$^{A}$ | VT044845-Gal4-DBD;<br>VT050660-p65AD | Bidaye et al. 2014 | Fig. 1j, 2f-g, 4g, 5h<br>Ext. Data Fig. 1c, 3b, 8b |
| MDN1$^{B}$ | VT049484-lexA-DBD;<br>VT050660-p65AD | Feng et al. 2020 | Fig. 3c-e, 4a-e<br>Ext. Data Fig. 4a,d-e, 5b, 6, 7b-f |
| DANs | R58E02-Gal4, TH-Gal4 | Kindly provided by Scott Waddell | Fig. 3e, 4a-e<br>Ext. Data Fig. 5b-g, 6, 7b-f |
| γ1 | MB320C-Gal4 | BDSC#68253 | Fig. 3c, 4h |
| γ2 | MB296B-Gal4 | BDSC#68308 | Fig. 3d |
| ChR2XXL$^{A}$ | UAS-ChR2-XXL | BDSC#58374 | Fig. 1c-d,f-i, 4g-h<br>Ext. Data Fig. 2a |
| ChR2XXL$^{B}$ | lexAop-ChR2-XXL | Kindly provided by Robert J. Kittel | Ext. Data Fig. 4d |
| Chrimson$^{A}$ | UAS-CsChrimson-mVenus | BDSC#55136<br>BDSC#55134 | Fig. 1e-g<br>Fig. 1a,j<br>Ext. Data Fig. 1a,c, 2b-c |
| Chrimson$^{B}$ | lexAop-CsChrimson-mVenus | Klapoetke et al. 2014 | Ext. Data Fig. 4e |
| Chrimson$^{C}$<br>GCaMP | lexAop-CsChrimson-tdtomato,<br>UAS-GCaMP6f | Perisse et al. 2016 | Fig. 3c-e, 4a-e<br>Ext. Data Fig. 5b-g, 6, 7b-f |
| Chrimson$^{D}$ | lexAop-ChrimsonR-mCherry | Kindly provided by Vivek Jayaraman | Ext. Data Fig. 4a-b |
| GtACR1 | UAS-GtACR1 | Kindly provided by Robert J. Kittel | Fig. 2f-g, 5h<br>Ext. Data Fig. 3b, 8b |
| CantonS | $CS_{gpu}$ | Kindly provided by Troy D. Zars | Ext. Data Fig. 8a |
| $w^{1118}$* | *white* null mutant | Hazelrigg et al. 1984 | Figure 1c,e-f,i, 2f-g<br>Ext. Data Fig. 2, 3b, 4d-e, 8b |
| pBDP* | Enhancerless Gal4 insert | BDSC#68384 | Figure 1j, 2f-g<br>Ext. Data Fig. 3b, 8b |

* Used to create heterozygous driver (Dri Ctrl) or effector controls (Eff Ctrl).
