## Supplemental Methods Table 2 for "Avoidance engages dopaminergic punishment in *Drosophila*"

| MBONs | FlyWire v783 IDs | LALs | FlyWire v783 IDs | MDNs (brain) | FlyWire v783 IDs |
|---|---|---|---|---|---|
| MBON30 | 720575940618008859 | LAL160,161 | 720575940626203394 | | 720575940616026939 |
| | 720575940637934308 | | 720575940617720577 | | 720575940631082808 |
| MBON35 | 720575940637902938 | | 720575940612624753 | | 720575940610236514 |
| | 720575940632943277 | | 720575940629547991 | | 720575940640331472 |
| MBON32 | 720575940609959637 | LAL042 | 720575940628016387 | | |
| | 720575940638526278 | | 720575940627802374 | **MDNs (VNC)** | **FANC IDs** |
| MBON27 | 720575940622979277 | LAL008 | 720575940639138382 | | 648518346475400628 |
| | 720575940618249797 | | 720575940643829704 | | 648518346474413506 |
| MBON26 | 720575940607155890 | LAL144a | 720575940619675200 | | 648518346501195096 |
| | 720575940629981440 | | 720575940620346417 | | 648518346481250703 |
| MBON31 | 720575940636992368 | LAL040 | 720575940617818708 | | |
| | 720575940644615716 | | 720575940622018217 | | |
| | | LAL171,172 | 720575940645716771 | | |
| | | | 720575940624891941 | | |
| | | | 720575940615651670 | | |
| | | | 720575940659019649 | | |
| | | LAL162 | 720575940637809326 | | |
| | | | 720575940630480459 | | |
| | | LAL051 | 720575940623067389 | | |
| | | | 720575940626694288 | | |
