## Supplemental Data Synapse Topology for "Avoidance engages dopaminergic punishment in *Drosophila*"

### Amin et al
### Supplemental Data Synapse Topology

X→MDN and MDN→X synaptic sites

LAL→MDN synaptic sites (Part 1)
LAL→MDN synaptic sites (Part 2)

MBON→LAL synaptic sites

### X→MDN and MDN→X synaptic sites

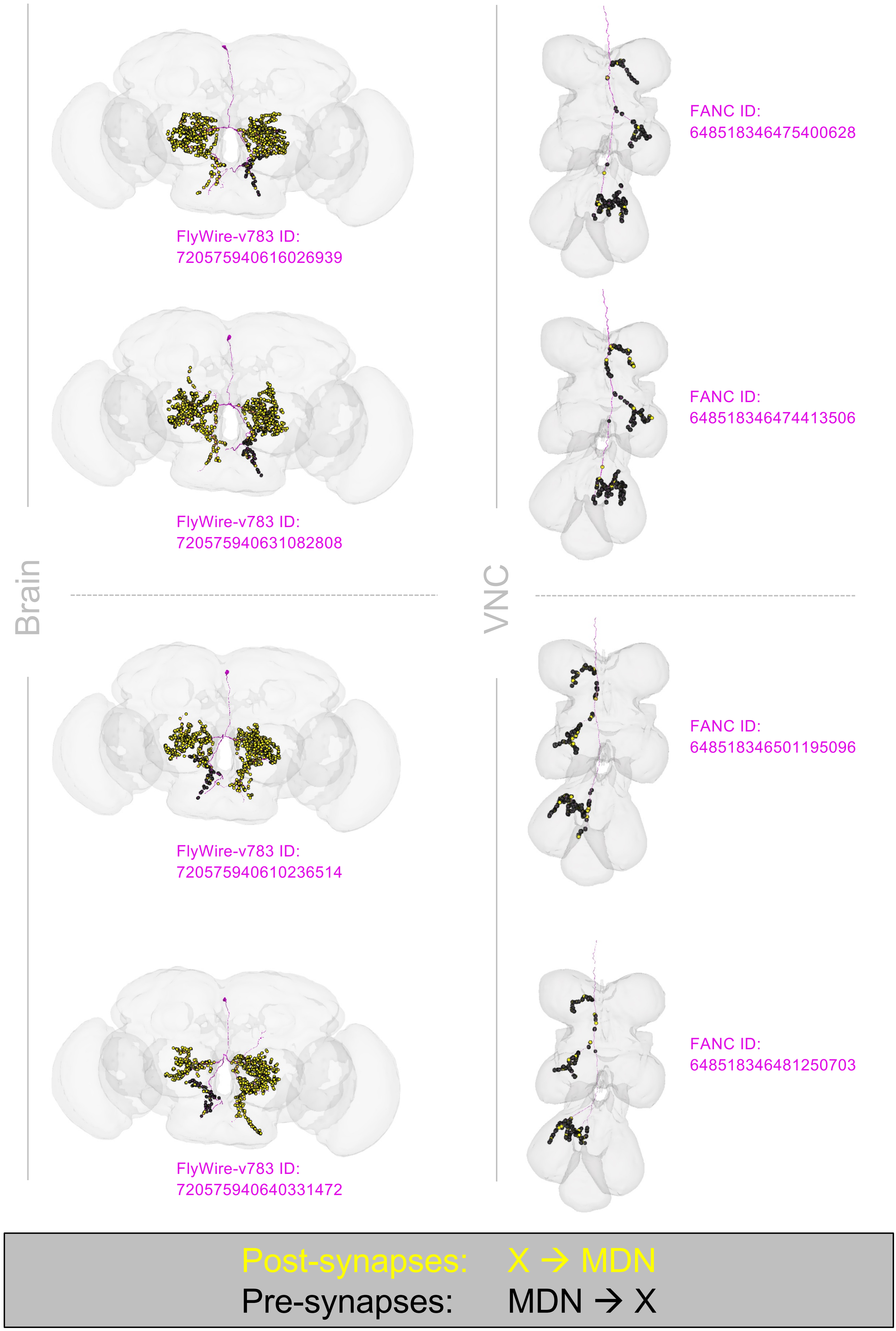

### LAL→MDN synaptic sites (Part 1)

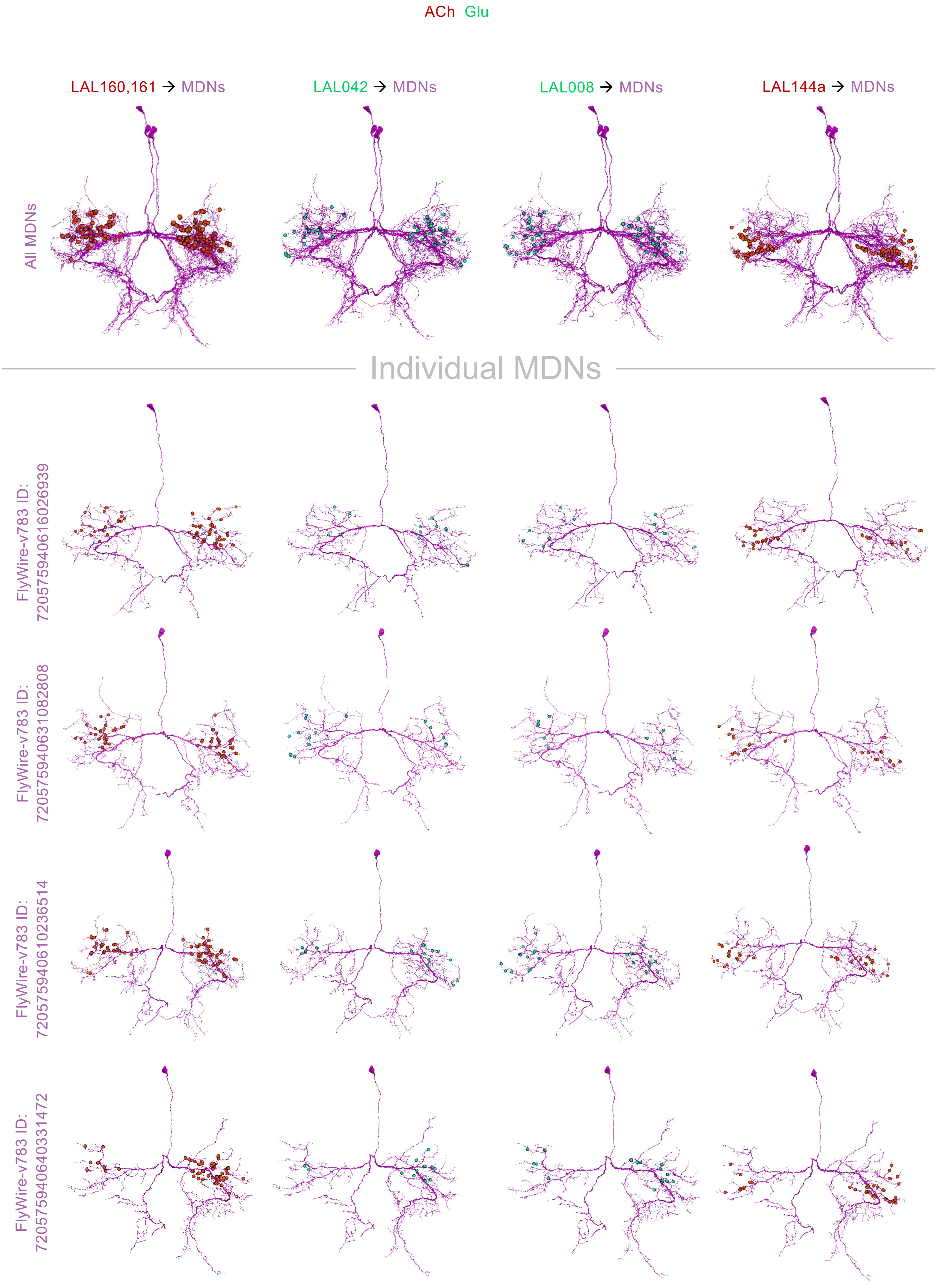

### LAL→MDN synaptic sites (Part 2)

ACh GABA Glu

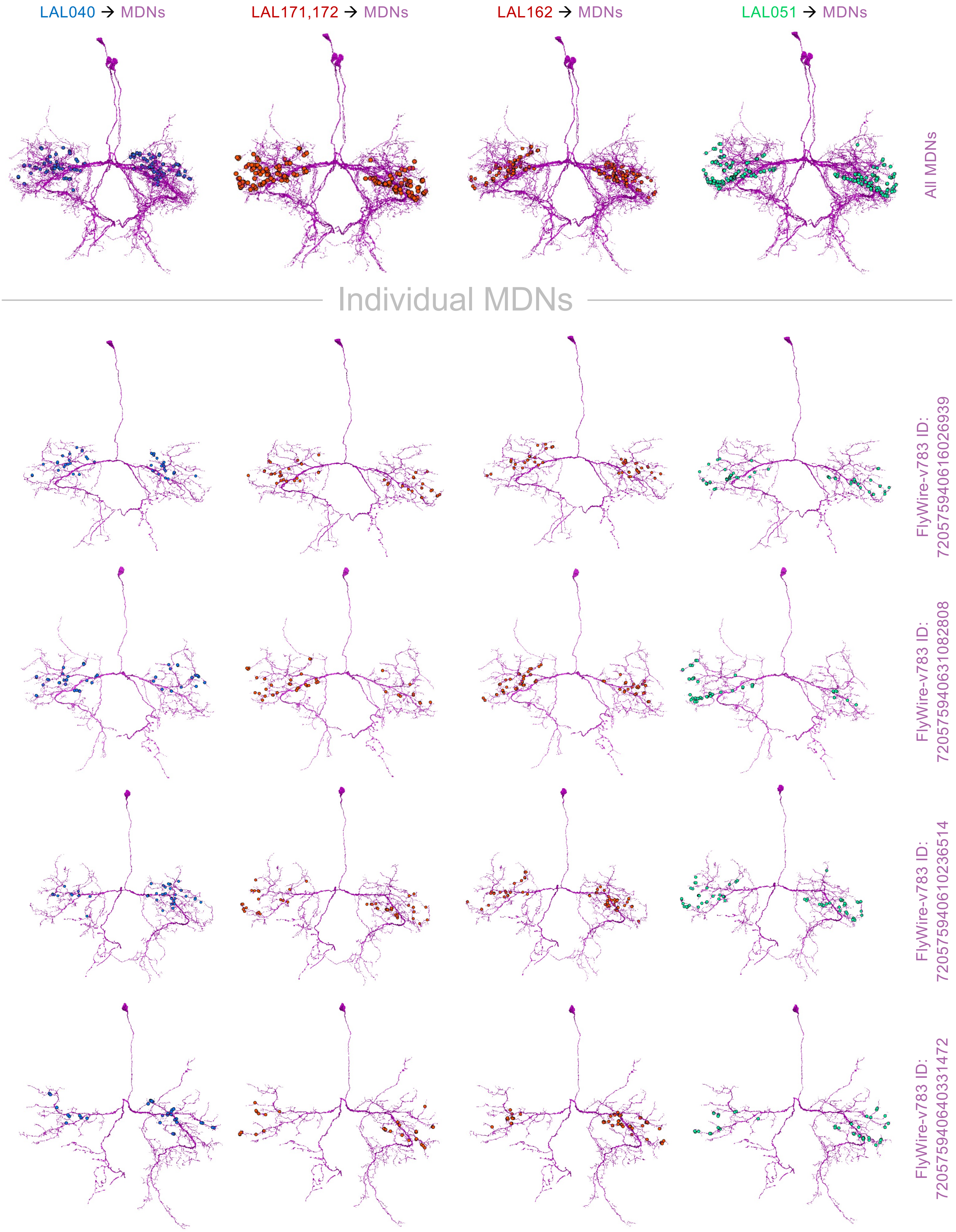

### MBON→LAL synaptic sites

ACh GABA Glu

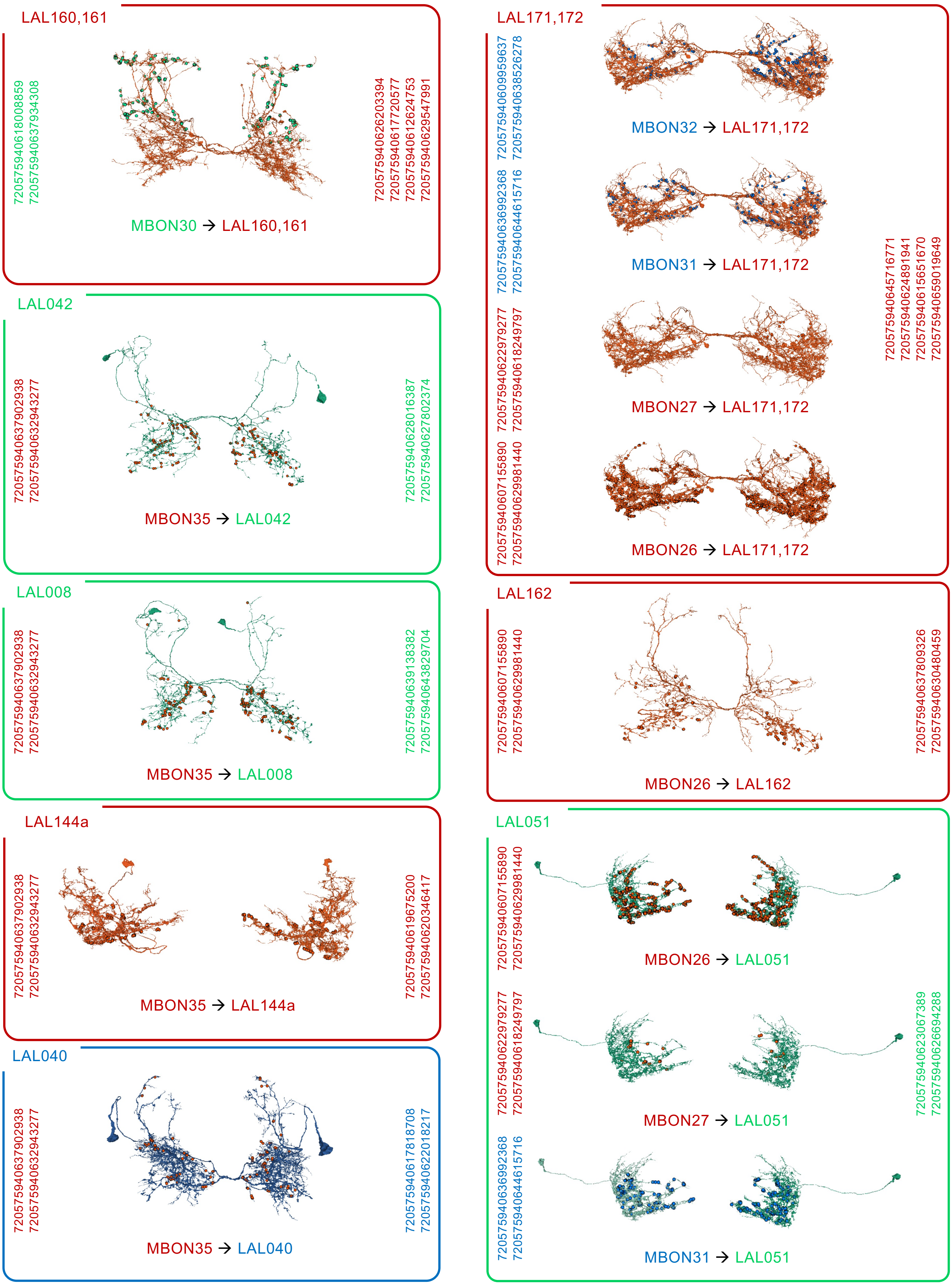
